# Cryo-EM study on the homo-oligomeric ring formation of yeast TRiC/CCT subunits reveals TRiC ring assembly mechanism

**DOI:** 10.1101/2021.02.24.432666

**Authors:** Caixuan Liu, Huping Wang, Mingliang Jin, Wenyu Han, Shutian Wang, Yanxing Wang, Fangfang Wang, Chun Su, Xiaoyu Hong, Qiaoyu Zhao, Yao Cong

**Affiliations:** State Key Laboratory of Molecular Biology, National Center for Protein Science Shanghai, Shanghai Institute of Biochemistry and Cell Biology, Center for Excellence in Molecular Cell Science, Chinese Academy of Sciences, Shanghai 200031, China; University of Chinese Academy of Sciences, Beijing 100049, China; The National Facility for Protein Science in Shanghai (NFPS), 201210 Shanghai, China; Shanghai Science Research Center, Chinese Academy of Sciences, Shanghai, China 201210

## Abstract

The complex eukaryotic chaperonin TRiC/CCT helps maintain cellular protein homeostasis, however, its assembly mechanism remains largely unknown. To address the subunit specificity in TRiC assembly, we express each of the individual yeast TRiC subunit in *E. coli*. Our cryo-EM structural study and biochemical analyses demonstrate that CCT1/2/6 can form TRiC-like homo-oligomeric double ring (HR) complex, however ATP-hydrolysis cannot trigger their ring closure; after deletion of the long N-terminal extension, CCT5 can form the closed double-ring structure; while CCT3/4/7/8 cannot form the HRs. It appears that CCT1 forms a HR in a unique spiral configuration, and ATP-hydrolysis can drive it to re-assemble with an inserted extra subunit-pair. Our data suggest that CCT5 could be the leading subunit in ATP-hydrolysis-driven TRiC ring closure. Moreover, we demonstrate that ADP is sufficient to trigger the assembly of the HRs and TRiC from the assembly intermediate micro-complex form. Our study reveals that through evolution, the more ancestral subunits may have evolved to take more responsibilities in TRiC ring assembly, and we propose a possible assembly mechanism of TRiC involving subunit-pair insertion. Collectively, our study gives hints on the structural basis of subunit specificity in TRiC assembly and cooperativity, beneficial for future TRiC-related therapeutic strategy development.

## Introduction

The eukaryotic group II chaperonin TRiC/CCT is the most complex chaperonin system, consisting of eight homologous but distinct subunits (CCT1 to CCT8 in yeast) arranged in a specific order, forming a double-ring configuration (Kalisman et al., 2012; Leitner et al., 2012; Wang et al., 2018; Zang et al., 2016; Zang et al., 2018). Each TRiC subunit consists of an ATP-binding equatorial domain and a substrate-binding apical domain linked by an intermediate domain. TRiC assists the folding of ∼10% cytosolic proteins including many key structural and regulatory proteins, such as actin, tubulin, cell cycle regulator CDC20, and cancer-related VHL tumor suppressor and STAT3 (Camasses et al., 2003; Dekker, 2010; Kasembeli et al., 2014; Llorca et al., 2000; Llorca et al., 1999; Trinidad et al., 2013). TRiC can assist the folding of such diversified substrates, mostly related to the structural asymmetry and specificity of TRiC subunits forming the complex assembly (Balchin et al., 2018; Jin et al., 2019; Leitner et al., 2012; Munoz et al., 2011; Plimpton et al., 2015; Roh et al., 2016; Zang et al., 2016).

Disfunction of TRiC has been closely related to cancer and neurogenerative diseases (Khabirova et al., 2014; Tam et al., 2009). Moreover, accumulating evidences have shown that the up-regulation or mutations of several individual subunits of TRiC are related to cancer, such as breast cancer (up-regulation of CCT1/CCT2/CCT5) (Guest et al., 2015; Ooe et al., 2007) and small cell lung cancer (up-regulation of CCT2) (Carr et al., 2017), without knowing the underling structural basis (Jin, 2019). Still, in the abovementioned diseases, it remains to be elucidated whether TRiC subunits act as a monomer, a homo-oligomeric complex, or a TRiC complex; if it functions as a monomer or a homo-oligomer, does it operate as a chaperone with ATPase activity or has other physiological functions. Interestingly, it has been reported that human CCT4 and CCT5 can form biologically active homo-oligomeric TRiC-like structure (Pereira et al., 2017; Sergeeva et al., 2013), and the synthetic CCT5 homo-oligomeric complex is able to inhibit mutant Huntingtin aggregation and promote the disassembly of mitotic checkpoint complex (MCC) in the same fashion as TRiC (Darrow et al., 2015; Kaisari et al., 2017; Shahmoradian et al., 2013). This lays down the necessity of studying the function of individual TRiC subunits, which may facilitate our understanding on the functionality of up-regulated TRiC subunit in the disease form.

Although TRiC has been discovered for decades, the biosynthetic pathway and assembly mechanism of intact TRiC remain poorly understood. It has been proposed that the micro-complex formed by several TRiC subunits might be the intermediate in the assembly pathway of intact TRiC, and TRiC assembly involves in the disassembly of TRiC in the lysate and then reassembly reaction whereby free subunits can assemble into the complex (Liou and Willison, 1997; Liou et al., 1998). Recently, it has been proposed that TRiC assembly relies on subunit exchange with some stable homo-oligomers, possibly CCT5, as base assembly units (Sergeeva et al., 2019).

To examine the homo-oligomeric ring-formation property of individual TRiC subunit so as to better understand the specificity of each subunit in TRiC assembly, we express and purify each of the eight yeast TRiC subunits in *E. coli*. Our systematic structural and biochemical analyses reveal that the individual CCT1/2/5/6 subunit can form homo-oligomeric ring-shaped (HR) structure like TRiC, and we resolve a series of cryo-EM structures of the TRiC-like HRs up to 3.2 Å resolution. Surprisingly, we find that CCT1 forms a HR in a unique spiral configuration with a seam, and ATP-hydrolysis can trigger it to re-assemble with one more inserted subunit-pair; CCT5 is more reactive responding to nucleotide and could be crucial in ATP-hydrolysis-driven TRiC ring closure. We find that the micro-complex is the assembly intermediate state of TRiC or TRiC-like ring, and ADP is sufficient to trigger their assembly from the micro-complex form. Our study shed light on the assembly mechanism of TRiC that may involve in subunit-pair insertion, and reveals that through evolution, the subunits from progenitor genes may be crucial in TRiC ring assembly.

## Results

### CCT1 can form a homo-oligomeric ring in a unique spiral configuration

To examine the ring formation property of individual TRiC subunit, we first expressed the CCT1 subunit of yeast TRiC in *E. coli*, and purified it without adding nucleotide (Fig. S1A). Our reference-free 2D classification and 3D reconstruction on the cryo-EM data both suggested that yeast CCT1 can form a homo-oligomeric double-ring structure (termed CCT1-HR) in an open conformation (Fig. 1A, S2A). Overall, the CCT1-HR complex consists of eight subunits per ring, and each individual subunit appears in the outward tilting conformation slanting away from the central cavity (Fig. 1A-B). Notably, the eight subunit pairs showed an unforeseen right-handed spiral configuration with a seam located between the highest subunit pair 8 and the lowest subunit pair 1, with a rise of ∼26 Å (Fig. 1A-C). Moreover, the seam subunit pairs 1 and 8 are less well resolved due to the dramatically reduced interaction interface area between them (Fig. S2C). Accordingly, the resolution of the map (6.1 Å) is not high due to the reduced inter-subunit interactions and intrinsic dynamics of the CCT1-HR complex (Fig. S2B-C).

**Fig. 1.**
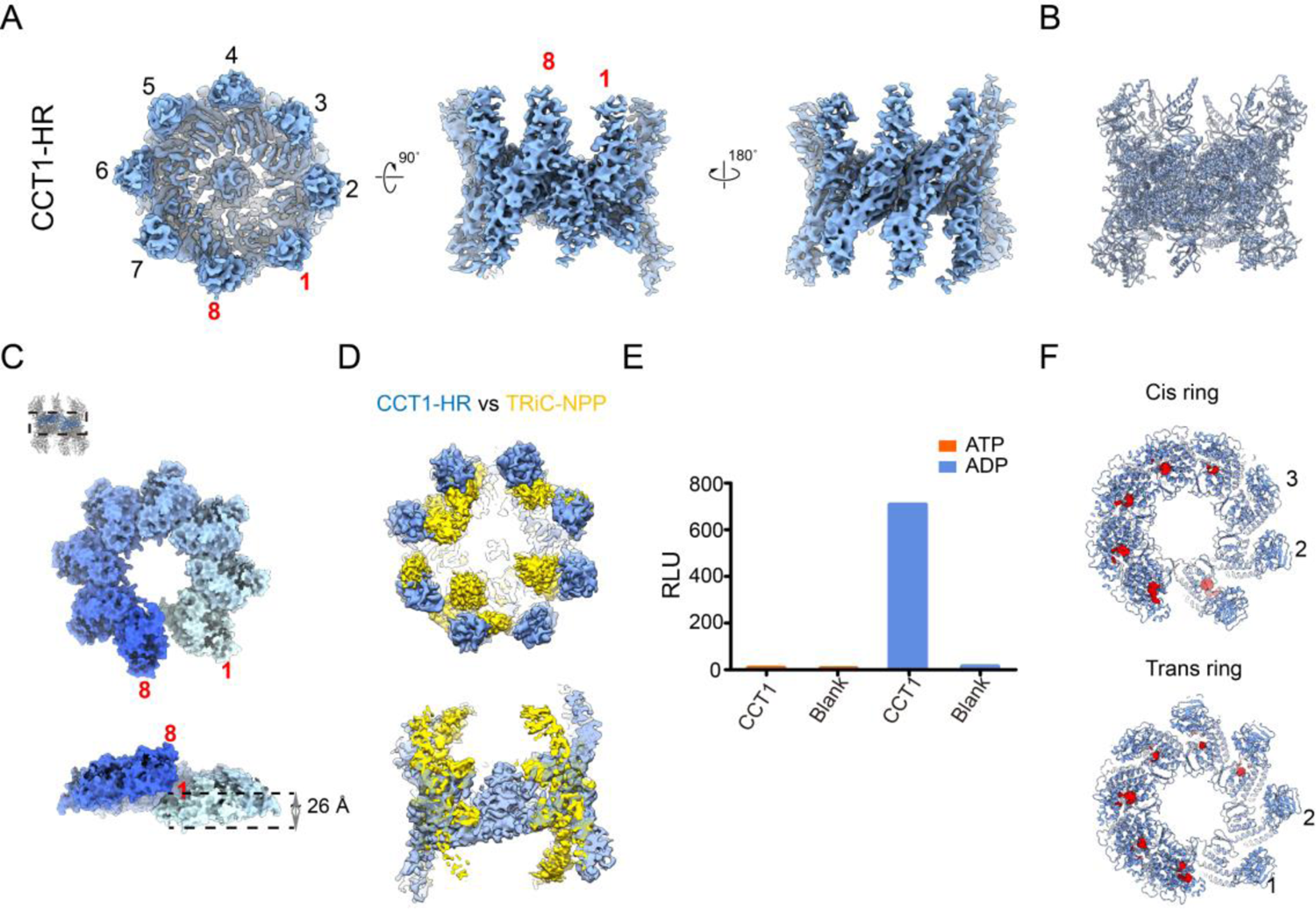
Cryo-EM structure of CCT1-HR revealing a unique spiral configuration. (A) Cryo-EM map of CCT1-HR, from left to right: top view (with the subunit pair numbered from the lowest 1 to the highest 8), and side views showing the seam/back of the seam regions. (B) The pseudoatomic model of CCT1-HR. (C) The spiral conformation of the E domain of CCT1-HR, with the rise between the lowest subunit pair 1 to the highest 8 being labeled. (D) Top and side views of the map overlay between CCT1-HR (cornflower blue) and yTRiC-NPP (yellow). CCT1-HR appears more open than yTRiC-NPP. (E) ATP/ADP ratio analysis of CCT1-HR purified without added nucleotide. The relative light unit (RLU) values were measured for the sample. (F) Nucleotide occupancy status of CCT1-HR, with the nucleotide density shown in red.

Interestingly, in the CCT1-HR structure, the conformation of CCT1 resembles that of the CCT1 in the WT yTRiC structure in the nucleotide partially preloaded state (TRiC-NPP) (Fig. S2D), consuming the most outward tilting conformation than all the other subunits (Zang et al., 2016). Consequently, the CCT1-HR complex appears more open than that of the TRiC-NPP (Fig. 1D) (Cong et al., 2012; Jin et al., 2019; Zang et al., 2016). Noteworthy, we observed another class (31.0% of the population) also presenting an overall eight-subunit-pair spiral configuration, but with the seam subunit-pair-8 missing simultaneously (Fig. S2E), indicating the two subunits may associate into a pair first then assemble into or leave the complex as a whole. We also speculate that this subunit-pair-8 missing configuration might represent an intermediate state during the double ring assembly or disassembly.

We then asked whether CCT1-HR has preloaded nucleotide from the environment of the cells as the WT yTRiC dose (Zang et al., 2016). Our ADP/ATP ratio analysis suggested that, although without added nucleotide during purification, CCT1-HR is in the ADP-bound state (Fig. 1E), which may come from the environment of *E. coli* cells. Substantiate to this, we found nucleotide densities in consecutive six subunits per ring in the CCT1-HR map, with subunits 2/3 in the cis-ring and 1/2 in the trans-ring un-occupied with nucleotide (Fig. 1F).

### ATP-hydrolysis cannot trigger the ring closure of CCT1-HR but can induce the ring to re-assemble

To investigate whether ATP-hydrolysis could trigger the ring closure of CCT1-HR, we incubated the complex with ATP-AlFx (the nucleotide analog mimicking the ATP-hydrolysis transition state) that can trigger the WT TRiC ring closure (Booth et al., 2008; Cong et al., 2010; Meyer et al., 2003). We resolved the cryo-EM map of the formed complex, termed CCT1-HR-ATP-AlFx, at 4.1-Å-resolution. Surprisingly, the complex remains in the open spiral conformation (Fig. 2A-B and S3A-B), indicating that ATP-hydrolysis cannot trigger the ring closure of CCT1-HR, instead, it can induce the ring to reassemble from 8 subunit pairs to 9 pairs, which to the best of our knowledge, has not been captured for TRiC related systems. Considering the weak interaction between the two seam subunit pairs and the co-existence of pair-8-missing configuration in CCT1-HR dataset (Fig. S2C, E), it is likely that the reassemble happens by squeezing in one more subunit-pair in the seam region induced by ATP hydrolysis. Also, the rise between the two seam pairs (9 and 1) increased dramatically from 26 Å to 70 Å (Fig. 2C), consequently the interaction between the seam subunit-pairs 9/1 is rather weak than that in the other non-seam neighboring subunit-pairs (Fig. S3C). Moreover, it appears that all the subunits have loaded nucleotides (Fig. 2D), and the subunits are in similar conformation (Fig. S3D). Corroborate to this, in the same dataset we obtained other two conformations with one or two seam subunit-pairs missing (Fig. S3F-G), suggesting that CCT1-HR-ATP-AlFx remains in a relatively dynamic subunit-pair reassemble/exchange status, similar to that of CCT1-HR. Together, it appears that ATP-hydrolysis is not sufficient to trigger the ring closure of CCT1-HR but results in reassemble of the complex with insertion of one more subunit-pair.

**Fig. 2.**
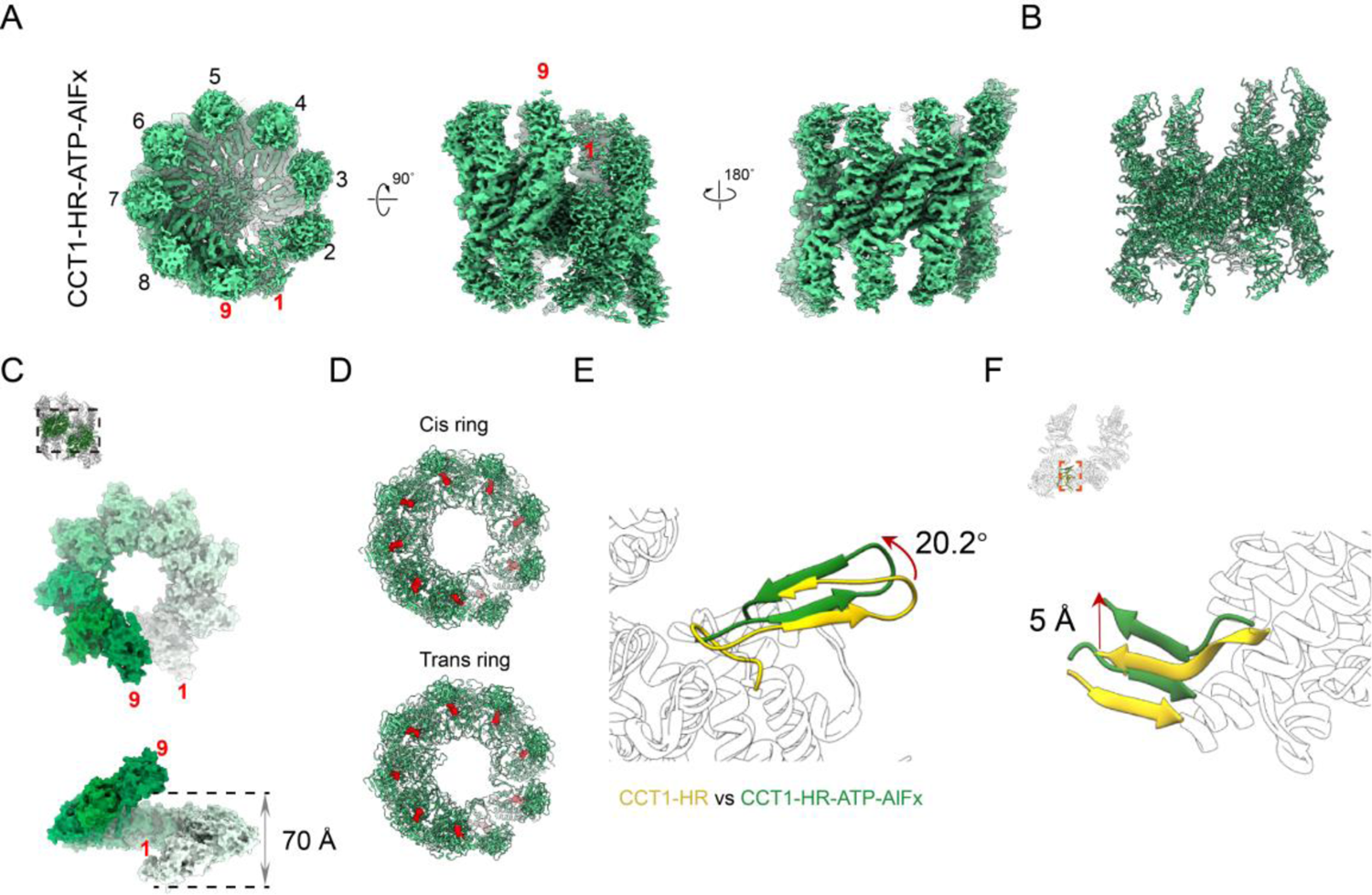
Cryo-EM structure of CCT1-HR-ATP-AlFx remaining in an open conformation. (A) The cryo-EM map of CCT1-HR-ATP-AlFx, from left to right: top view (with the subunit pair numbered from the lowest 1 to the highest 9), and side views from the seam/the back of the seam regions. (B) The pseudoatomic model of CCT1-HR-ATP-AlFx. (C) The spiral conformation of the E domain of CCT1-HR-ATP-AlFx, with the rise between the lowest subunit pair 1 to the highest 9 being labeled. (D) Nucleotide occupancy status of CCT1-HR-ATP-AlFx. (E and F) Model alignment for the neighboring subunits between CCT1-HR (gold) and CCT1-HR-ATP-AlFx (forest green), aligned by the lift subunit. Showing is that ATP-hydrolysis could lead to an obvious upward lift of ∼20.2° in the stem loop of the left subunit (E), coupled with a simultaneous lift of ∼5 Å of the N/C termini of the right neighboring subunit (F).

We then examined the structural mechanism of the rise increasement induced by ATP-hydrolysis and the resulted ring reassembly in CCT1-HR-ATP-AlFx. Subunit model alignment between CCT1-HR and CCT1-HR-ATP-AlFx suggested that ATP-hydrolysis could lead to an obvious ∼20.2° upward lift in the stem loop region (Fig. 2E and S3E), and the immediately coupled N/C termini of the right neighboring subunit showed a simultaneous lift of ∼5 Å to maintain the association between the two neighboring subunits (Fig. 2F). This way, the ATP-hydrolysis induced conformational changes originated in the stem loop region could be propagated to the neighboring subunits inducing the spiral rise enlargement (Fig. 2C), which could potentially break the originally weak interaction between the seam subunit-pairs and lead to the re-assemble and insertion of one more subunit-pair in the seam region, forming the 9-meric ring complex.

### ATP-hydrolysis can induce the CCT2 A/I domain unbending but not sufficient to trigger the CCT2-HR ring closure

We also expressed yeast CCT2 subunit in *E. coli* and purified it without adding nucleotide throughout the purification process (Fig. S1A). Our cryo-EM reconstruction at 4.5-Å-resolution showed that CCT2 can as well form a homo-oligomeric double-ring (CCT2-HR) structure with eight subunits per ring (Fig. 3A and S4A and C). Overall, the A/I domains together adopt bended conformation and are less well resolved, indicating an intrinsic dynamic nature of the two domains (Fig. 3A, B). The A/I domains tend to contact with the left neighboring subunit, similar to that of the CCT2 subunit in the WT yTRiC-NPP structure (Zang et al., 2016), forming four subunit-pairs in the better resolved cis-ring (Fig. 3A). In addition, the ADP/ATP ratio assay suggested that CCT2-HR is also in the ADP bound state although no added nucleotide in the purification process (Fig. 3C).

**Fig. 3.**
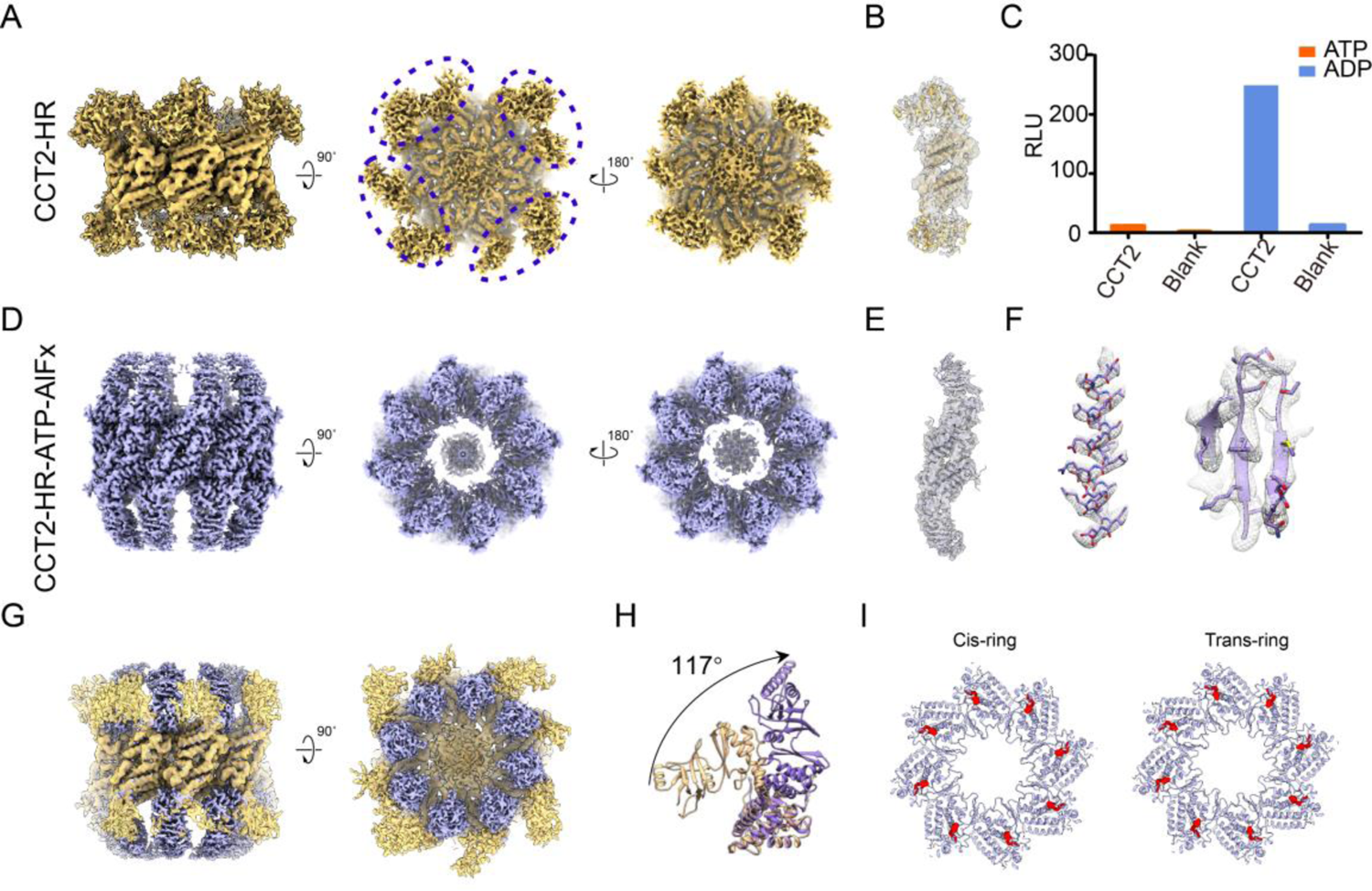
Cryo-EM structures of CCT2-HR and CCT2-HR-ATP-AlFx. (A) The cryo-EM map of CCT2-HR. From left to right: side view and two end-on views. (B) Map and model fitting for a single subunit of CCT2-HR. (C) ATP/ADP ratio analysis of CCT2-HR purified without added nucleotides. (D) The cryo-EM map of CCT2-HR-ATP-AlFx, showing is the side view and two end-on views. (E-F) Good match between the map and model of a single subunit of CCT2-HR-ATP-AlFx (E), and representative high-resolution structural features. (G-H) Overlay of the CCT2-HR (in gold) and CCT2-HR-ATP-AlFx (in purple) maps (G) and models of a single-subunit pair (H), showing that ATP-hydrolysis can trigger the A/I domain unbending of CCT2 with an angle of 117°. (I) Nucleotide occupancy status of CCT2-HR-ATP-AlFx.

Furthermore, we resolved a CCT2-HR-ATP-AlFx map at 3.2-Å-resolution (Fig. 3D-F, S4B-D). Of note, the complex remains in an open conformation, although exhibiting an ATP-hydrolysis induced large unbending of ∼117° in the A/I domains of CCT2 (Fig. 3G-H), consuming an origination similar to that of the CCT2 in the yTRiC-AMP-PNP structure (Fig. S4E) (Zang et al., 2016). Correlatively, we detected nucleotide densities in all subunits in this stabilized structure (Fig. 3I). Altogether, it appears that ATP-hydrolysis can induce the A/I domain unbending of CCT2 consuming the ATP-binding conformation but not sufficient to trigger CCT2-HR ring closure.

### CCT5 needs the presence of nucleotide to form the ring and its long N-terminus could hinder the double ring formation

For yeast CCT5 subunit, we found that without added nucleotide during purification, it cannot form the ring-shaped complex, but rather much smaller micro-complexes (Fig. S8A, left panel). However, adding ATP (1 mM) in the purification process can induce the formation of the CCT5 homo-oligomer ring with eight subunits and in the closed state (Fig. S5A). We then resolved the cryo-EM map of the CCT5 homo-oligomeric ring at 4.0-Å-resolution (Fig. 4A-B, S5A, C-D), which, surprisingly, is in an unforeseen single-ring configuration (termed CCT5-HSR-ATP). Our structural analysis revealed that all the eight subunits have bound nucleotide in CCT5-HSR-ATP (Fig. 4C), and its ring appears more compact with the A-domain helical-protrusion of CCT5 undergoing a ∼14.2° downward tilt comparing with that of CCT5 in the closed state of WT yTRiC complex (Fig. 4G, PDB: 6KS6) (Jin et al., 2019).

**Fig. 4.**
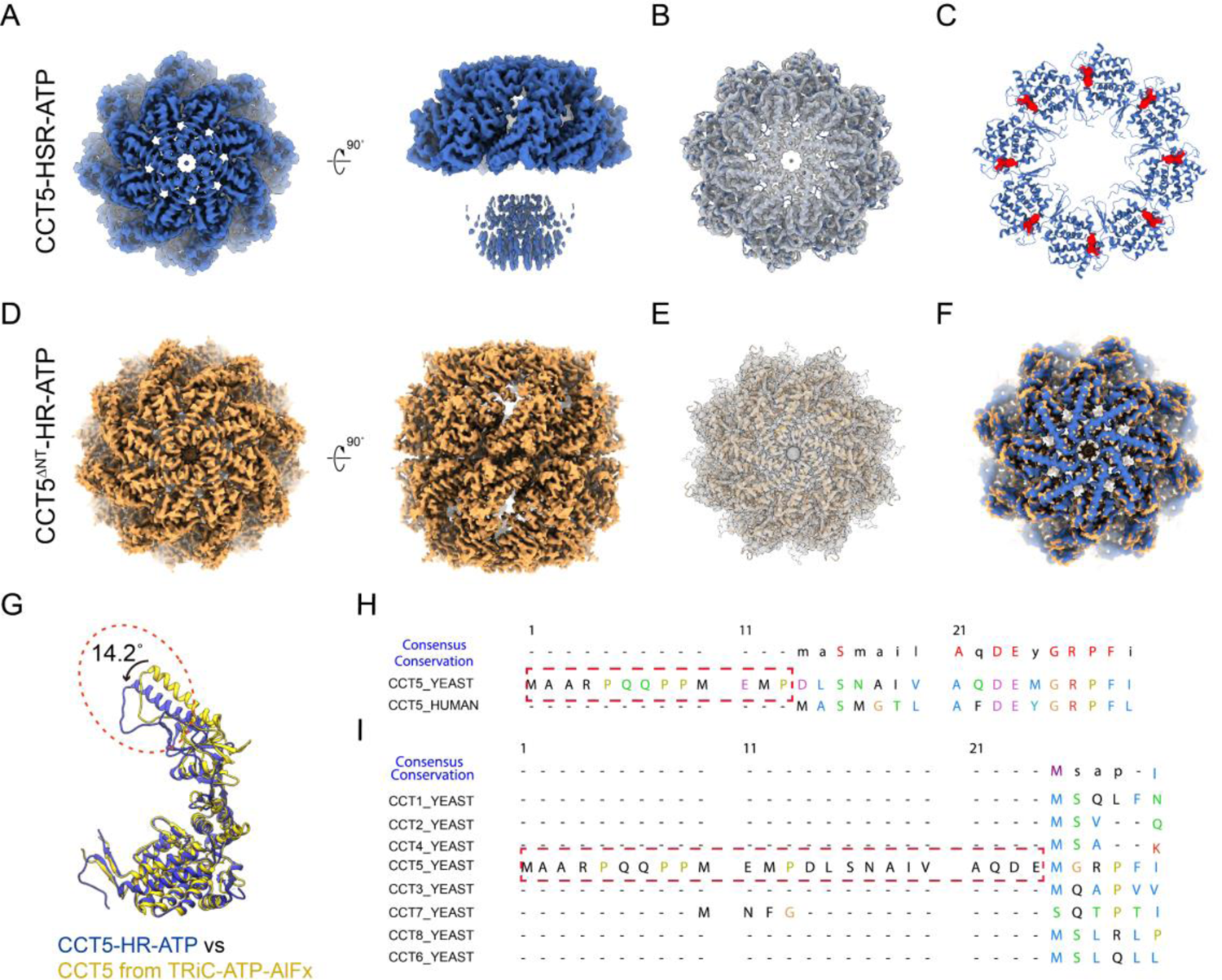
Cryo-EM structures of CCT5-HSR-ATP and CCT5^ΔNT^-HR-ATP and the extremely long N-terminus of yCCT5. (A) The cryo-EM map of CCT5-HSR-ATP, showing is the top and side views. (B) Good match between the map (transparent gray) and the pseudoatomic model (blue) of CCT5-HSR-ATP. (C) The nucleotide status of CCT5-HSR-ATP. (D) The cryo-EM map of CCT5^ΔNT^-HR-ATP, showing is top and side views. (E) Good match between the map and pseudoatomic model of CCT5^ΔNT^-HR-ATP. (F) Overlay of the CCT5-HSR-ATP and CCT5^ΔNT^-HR-ATP maps, indicating no obvious conformational variation between them. (G) Aligned model of the CCT5 subunit from CCT5-HSR-ATP (blue) and TRiC-ATP-AlFx (PDB: 6KS6, yellow), showing that the CCT5 A-domain helical-protrusion undergoes a ∼14.2° inward tilt from TRiC-ATP-AlFx to the CCT5-HSR-ATP. (H) Sequence alignment between yCCT5 and hCCT5, showing is the N-terminal region. (I) Sequence alignment of the eight subunits of yTRiC indicating the relative long N-terminus of yCCT5.

Further sequence alignment showed that the N-terminus of yeast CCT5 is 13 residues longer than its human counterpart (Fig. 4H); and among the eight subunits of yeast TRiC, CCT5 has the longest N-terminal extension with an extra ∼24 amino acids (Fig. 4I). We then postulate that the long N-tails from the eight CCT5 subunits forming the CCT5-HR-ATP ring, extending toward the two-ring interface (Fig. 4A), could group together physically blocking the double ring association. This also explain why human CCT5 can form double ring structure while yeast CCT5 cannot (Sergeeva et al., 2013). To validate our hypothesis, we truncated the N-terminal 24 amino acids of CCT5 (termed CCT5^ΔNT^). We found that CCT5^ΔNT^ can naturally form closed double ring structure with or without added nucleotide (Fig. S5F), and it has been documented that the human CCT5 can form double ring structure in the presence of ATP (Sergeeva et al., 2013). Moreover, in the absence of added nucleotide CCT5^ΔNT^ is not stable enough to sustain the cryo vitrification process, thus we only determined the cryo-EM map of CCT5^ΔNT^-HR-ATP at 3.7-Å-resolution (Fig. 4D-E, Fig. S5B-C, E), which is in closed double-ring configuration, with the ring conformation resembles that of CCT5-HSR-ATP (Fig. 4F). Collectively, our data demonstrate that the long N-tail of yeast CCT5 could block the double ring association in CCT5-HSR-ATP, however after deletion of its long N-tail, it can form the closed double-ring structure.

### CCT6 can also form the HR but ATP-hydrolysis cannot trigger the ring closure

Here we also showed that CCT6 can form a double-ring structure (CCT6-HR) without addition of ATP during purification (Fig, S1A, Fig. 5A). Noticing the CCT6-HR exhibited obvious preferred orientation problem, we then adopted the stage tilt strategy to collect additional tilted data (Fig. S6A, Table1) (Tan et al., 2017), which has been shown to help to overcome the preferred orientation problem in cryo-EM study of various complexes (Xu et al., 2020; Zheng et al., 2021). Noteworthy, CCT6-HR displays compositional heterogeneity, and we obtained an 8- and a 9-meric double ring structure both at open conformation (Fig. 5A, Fig. S6A-D). In the better resolve 8-meric CCT6-HR structure (at 5.9-Å-resolution), the subunit conformation resembles that of the CCT6 in WT TRiC-NPP complex (PDB: 5GW4, Fig. S6E) (Zang et al., 2016). In addition, we can detect nucleotide density in all subunits of CCT6-HR (Fig. 5B), and the ADP/ATP ratio assay suggested that although no added nucleotide, CCT6-HR is also in the ADP bound state, similar to that of CCT1 and CCT2 (Fig. 5C).

**Fig. 5.**
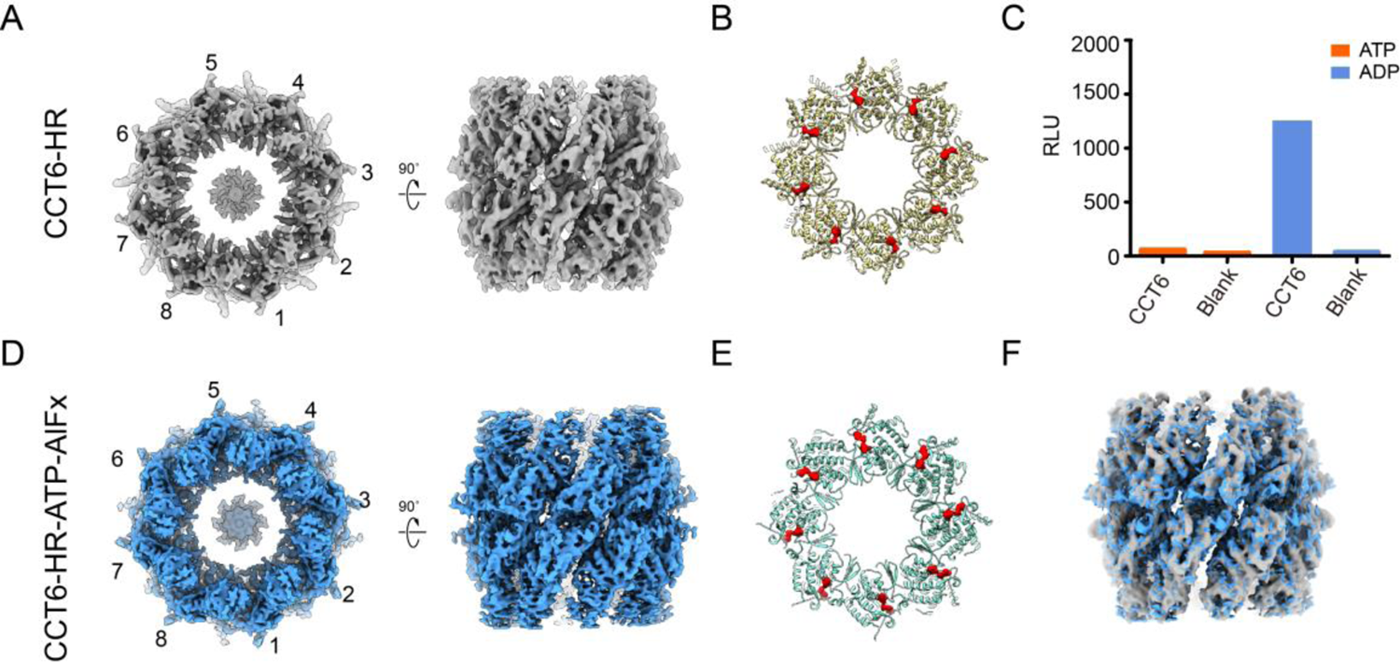
Cryo-EM structures of CCT6-HR and CCT6-HR-ATP-AlFx. (A) The cryo-EM map of CCT6-HR, showing is the top and side views. (B) The nucleotide occupancy statuses of CCT6-HR. (C) ATP/ADP ratio analysis of CCT6-HR purified without added nucleotides. (D) The cryo-EM map of CCT6-HR-ATP-AlFx. (E) The nucleotide occupancy statuses of CCT6-HR-ATP-AlFx. (F) Overlay of the CCT6-HR and CCT6-HR-ATP-AlFx maps, indicating no conformational difference between them.

Furthermore, we resolved the cryo-EM maps of CCT6-HR-ATP-AlFx, including an 8-meric structure at 4.2-Å-resolution and a 9-meric ring structure (Fig. 5D-E, S7), both remaining open. Interestingly, there is no obvious conformational changes between the 8-meric CCT6-HR-ATP-AlFx and CCT6-HR structures (Fig. 5F), suggesting that ATP-hydrolysis cannot trigger CCT6-HR ring closure.

### The yeast CCT3, CCT4, CCT7, and CCT8 subunit cannot form the ring-like structure

We also individually expressed yeast CCT3, CCT4, CCT7, and CCT8 subunits in *E. coli* following similar procedure described above. Although can be expressed, each of the subunits cannot form homo-oligomeric ring-shaped structure with or without added ATP. Instead, each of them can form flaky structure of unknown oligomeric numbers with loaded ADP most likely from the environment (Fig. S1B-I).

### Nucleotide is essential in the ring assembly of TRiC or TRiC-like HRs

We found that for the CCT5 micro-complex, adding ATP or ADP could trigger it to assemble into ring-shaped complex (Fig. S8A). However, for the CCT5 monomer, it remains in the monomeric form regardless of the presence or absence of ATP or ADP (Fig. S8B). Therefore, for CCT5, the micro-complex could be an essential assembly intermediate, from where adding nucleotide could trigger the ring assembly.

We then examined whether WT yTRiC has similar property. We first found that by mixing yTRiC with a buffer containing higher salt concentration but no glycerol at room temperature for 4 hours (Method), the complex can be disassociated into micro-complex (Fig. S8C, right panel). Interestingly, by adding ATP or ADP, it appears that the disassociated TRiC can re-assemble into ring-shaped structure (Fig. S8D) either in the closed (with ATP) or open (with ADP) conformation, implying the re-assembled TRiC remains its ATPase activity (Cong et al., 2012; Jin et al., 2019). Altogether, the micro-complex of TRiC is an essential assembly intermediate, from where, adding nucleotide could trigger complete TRiC ring assembly. Still, there might be unknown assembly factors for TRiC micro-complex formation from monomeric formation.

Here we showed that for CCT1/2/6-HRs, they remain bind ADP from the environment of the cells. It has been documented that the strictly conserved P-Loop (GDGTT motif), located in the E domain of group II chaperonins, is heavily involved in nucleotide binding (Amit et al., 2010). To further address the role of nucleotide in TRiC ring assembly, we replaced the P-Loop (GDGTT) with AAAAA in CCT1/2/5/6, respectively (Fig. 6A). Our ATP/ADP ratio assay revealed that these P-loop mutants cannot bound nucleotide anymore (Fig. 6B), in line with a previous report (Reissmann et al., 2012). Consequently, each of these mutants can only form micro-complex instead of the HR regardless of the presence or absence of ATP (Fig. 6C). Collectively, our data demonstrate the essential role of nucleotide-binding in the assembly of TRiC or TRiC-like HR complex.

**Fig. 6.**
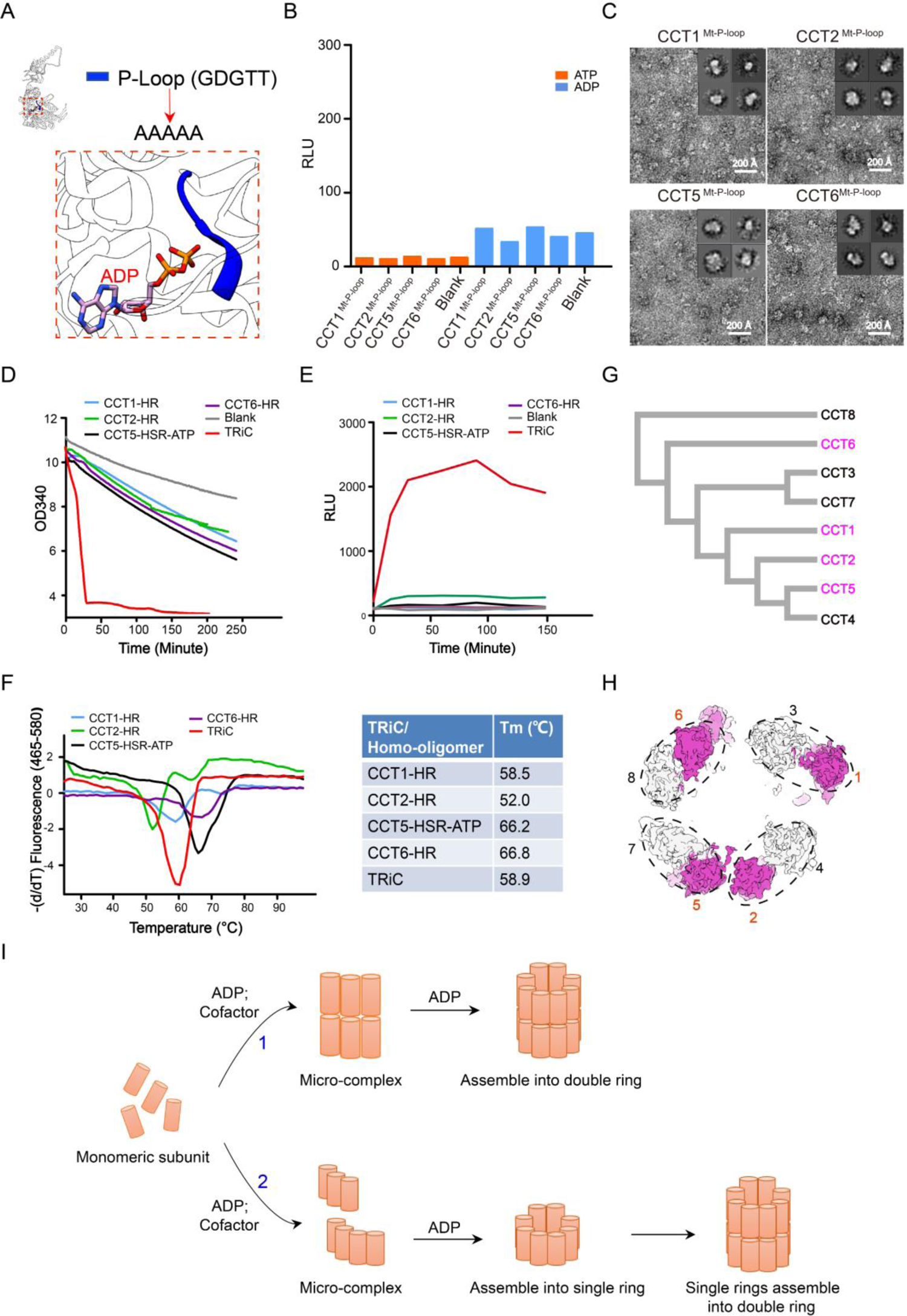
The importance of nucleotide-binding in the formation of TRiC-like HRs, functionalities of the TRiC-like HRs, and the possible assembly modes of intact TRiC. (A) The P-loop (GDGTT-motif, in blue) location of TRiC. Here we mutate the P-loop into AAAAA to investigate the role of nucleotide-binding in the formation of TRiC-like HRs. (B) ADP/ATP ratio assay for the P-loop mutants of CCT1/2/5/6. (C) The representative NS-EM micrograph and reference-free 2D class averages of the P-loop mutants of CCT1/2/5/6 purified in the presence of 1 mM ATP, indicating that the P-loop mutants cannot form HRs but micro-complexes. (D and E) ATPase activity assay (D) and Luciferase folding assay (E) of CCT1/2/5/6 HRs and yTRiC, showing that yeast CCT1/2/5/6 HRs have lower ATPase activity and rather weak substrate processing ability. (F) Thermal ability analysis of the HRs, in which the peak location in the X-axis correlates to the complex disassociation temperature. (G) Phylogenetic analysis on the eight subunits of yTRiC, suggesting that CCT4/5 are the progenitor genes, CCT2 and CCT1 are relatively close to CCT4/5, and CCT7/3/6/8 originated by duplication of an CCT4/5/2/1 genes. The four subunits which can form HRs are shown in violet red. (H) The four subunit pairs in TRiC-AMP-PNP (PDB: 5GW5) are highlighted by dotted black ellipsoid. The four HR-forming subunits are colored in violet red. (I) Two possible assembly modes of intact TRiC complex.

### Functionalities of the TRiC-like HRs

Furthermore, our ATPase activity and luciferase-folding ability assays showed that the HRs formed by yTRiC individual subunit have ATPase activity although obviously weaker than that of WT yTRiC (Fig. 6D), and they displayed rather low or even negligible luciferase-folding activity compared with that of yTRiC (Fig. 6E). These data suggest that for yeast TRiC, only when the diversified subunits assemble together in certain order forming contact TRiC, they can transform into a highly allosterically coordinated machine that can efficiently consume ATP to drive its folding cycle for productive substrate folding (Lopez et al., 2017; Reissmann et al., 2012).

It has been reported that the ATP hydrolysis activity and peptide refolding ability of recombinant chaperonin thermosomes of different species are quite different. For example, the recombinant thermosomes from *S. shibatae* and *S. solfataricus* show high ATP hydrolysis activity and facilitate refolding of unfolded proteins (Guagliardi et al., 1994; Kagawa et al., 1995); whereas the recombinant ones from *S. acidocaldarius*, *S. tokodaii* or *Acidianus tengchongensis strain S5T* exhibited trace ATP hydrolysis activity and peptide refolding ability in vitro (Minuth et al., 1999; Wang et al., 2010; Yoshida et al., 1998), but some of them can prevent mesophilic substrates from irreversible thermal aggregation (Wang et al., 2010). Therefore, although the HRs formed by yTRiC subunits CCT1/2/5/6 show lower ATPase activity and weak luciferase folding ability (Fig. 6D-E), they may have other functionalities such as preventing substrate aggregation.

In addition, the stability measurement through the thermal shift assay (Fig. 6F) showed that CCT6-HR has the highest Tm value, indicating an obviously high-stability of the complex, which is consistent with our previous observation that CCT6 is usually more stable and better resolved than the other subunits in the open-state TRiC (Cong et al., 2012; Zang et al., 2016). The CCT5-HSR-ATP in the closed-ring configuration is also highly stable, probably due to increased intra-ring subunit interactions. CCT2-HR has the lowest Tm value, in line with its structural feature that the neighboring subunits show least contacts due to the A/I-domain bending (Fig. 3A-B). Interestingly, although CCT1-HR is fairly dynamic in the seam, it shows comparable Tm value with WT TRiC.

## Discussion

Investigation on the ring-formation property of individual TRiC subunit could facilitate our understanding on the specificity of each subunit in TRiC assembly and coordination. Unlike the previous study focused on the human CCT4 and CCT5 subunits (Sergeeva et al., 2013), here we independently expressed and purified the eight subunits of yTRiC in *E. coli*. Our systematic cryo-EM structural study combined with biochemical and functional analyses revealed that yeast CCT1/2/6 can form TRiC-like homo-oligomeric double-ring structures; CCT5 with an extremely long N-tail can form a closed single-ring structure in the presence of ATP; while yeast CCT3/4/7/8 cannot form the HR structure regardless of the presence or absence of ATP. Interestingly, all the TRiC-like HRs show distinct structural and functional specificities: (1) CCT1-HR exhibits a spiral configuration with a seam, and ATP-hydrolysis can induce it to re-assemble from 8- to 9-meric ring configuration with an inserted subunit-pair; (2) the originally bent A/I-domains of individual subunit in CCT2-HR can be unbent induced by ATP-hydrolysis, but this is not sufficient to trigger the ring closure of the complex; (3) for yeast CCT5, the extremely long N-tail could block the double-ring formation, however, after N-tail truncation CCT5^ΔNT^ can form a closed double-ring structure regardless of the nucleotide state; (4) CCT6-HR shows compositional heterogeneity, and the presence of nucleotide cannot close the ring. Our study reveals specific characteristic of individual yTRiC subunit in the TRiC-like HR assembly as well as distinct structural and functional specificities of these formed HRs, facilitating our understanding of the subunit specialty in the assembly and allosteric cooperativity of WT TRiC.

Our finding that CCT1/2/5/6 can form HRs while the remaining CCT3/4/7/8 cannot is comparable with that of chaperonin *Thermoplasma acidophilum* (*Ta*) thermosome, in which only the α subunit could form the HR while β subunit cannot (Shoemark et al., 2018). Similarly, for human proteasome 20S, only the α7 subunit could self-assemble into a double-ring tetradecamer (Gerards et al., 1997; Kozai et al., 2017). Our further phylogenetic analysis on the yTRiC subunits showed that among the four HR-forming subunits CCT1/2/5/6, three of them (CCT1/2/5) are the progenitor genes (Fig. 6G) (Mukherjee et al.; Pereira et al., 2017). This is comparable with the *S. shibatae* TF55 thermosome that only the more primitive α and β subunits could form the HRs while γ subunit cannot (Kagawa et al., 2003). Moreover, we have previously revealed a tetramer-of-dimer pattern among the eight subunits in the open conformation of TRiC (Cong et al., 2012; Zang et al., 2016). It appears that within each dimer there is a HR-forming subunit (Fig. 6H), indicating that the intervallic arrangement of the four HR-forming subunit is sufficient to accommodate the remaining subunits for the WT TRiC ring formation, thus presenting a tetramer-of-dimer pattern. Collectively, it appears that through evolution, the subunits from the progenitor genes may play more essential roles in TRiC ring assembly.

### CCT5 could be the leading subunit in ATP-hydrolysis-driven TRiC ring closure

Interestingly, our data show that in the recombinant CCT1-, CCT2-, and CCT6-HR complexes, the individual CCT1/2/6 subunit remains in their original conformation as in the WT yTRiC-NPP complex (Figs. S2D, S4E, and S6E), and ATP-hydrolysis is not sufficient to trigger their HR ring closure, but can drive CCT2-HR to the ATP-binding conformation. In contrast, after deletion of the long N-tail, CCT5^ΔNT^ can naturally form closed double ring structure with or without added nucleotide (Figs. 4A, S5F). Our finding is substantiated by a previous biochemical study suggesting that among all viable TRiC-subunit HYD (key for ATP-hydrolysis) mutants, a notable exception was the TRiC-CCT5 HYD mutant, in which the lid closure was substantially impaired (Reissmann et al., 2012). Collectively, CCT5 could be the leading subunit in responding to ATP-hydrolysis to eventually drive the TRiC ring closure, while CCT1/2/6 and most likely the other subunits may need to be allosterically coordinated to involve in TRiC ring closure.

### ADP is sufficient to trigger TRiC or TRiC-like ring assembly from the intermediate micro-complex form

It has been investigated on the role of nucleotide in driving TRiC ring conformational transition (Booth et al., 2008; Cong et al., 2012; Meyer et al., 2003; Reissmann et al., 2012). Here our biochemical and structural data further demonstrated that nucleotide is essential for the homo-oligomeric ring assembly: (1) for CCT1/2/6 HR and CCT3/4/7/8 micro-complexes, they all have bound ADP from the environment of cells; (2) for P-loop mutated CCT1/2/5/6 that unable to bind nucleotide anymore, they cannot form ring-shaped complex but only micro-complexes; (3) the micro-complex of CCT5 or disassociated yTRiC can assemble into ring-shaped structure in the presence of ATP or ADP. This further suggests that ADP-binding is sufficient to trigger TRiC or TRiC-like ring assembly from the micro-complex form, in line with a previous report suggesting that nucleotide binding is necessary and sufficient to induce thermosome assembly (Zhang et al., 2013). Combined with our prior observation that in WT yTRiC, CCT3/6/8 have preloaded ADP from the environment (Jin et al., 2019; Zang et al., 2016), it appears that ADP could be sufficient for the assembly and stabilization of the TRiC complex, while ATP could provide energy to drive the TRiC ring closure for substrate folding.

### The possible mode of TRiC complex assembly

It has been a long-standing question regarding the mechanism of TRiC complex assembly. Here we found that triggered by ATP-hydrolysis, yeast CCT1-HR can re-assemble from the 8- to 9-meric ring by squeezing in one more vertical subunit-pair in the weakly associated seam region, and also captured the assembly intermediate state with missing subunit pairs (Fig. S3F-G). It is possible that TRiC assembly relies on subunit-pair exchange with some homo-oligomers, possibly CCT1-HR in yeast, as base assembly unit. Overall, based on our study, we postulate a subunit-pair association mechanism for the *ab initio* TRiC assembly: two matching TRiC subunits oppositely stand on top of each other in their E domains forming a stable subunit pair, and such subunit pairs gradually associate into micro-complexes, which in the presence of ADP could then assemble into the complete double-ring TRiC complex following certain order (Fig. 6I-1). Still, we cannot rule out the possibility that TRiC subunits can first assemble into micro-complexes, then into single ring in the presence of ADP, eventually two single rings associate into a double-ring TRiC complex (Fig. 6I-2). Considering that for yCCT5, after deletion of the long N-tail it can still form the closed double-ring structure, we prefer the assembly mechanism 1 (Fig. 6G-1).

In summary, our studies provide a through picture on the homo-oligomeric ring formation property of yTRiC individual subunit and the characteristics of those formed TRiC-like HRs. We show the micro-complex of TRiC or TRiC-like HRs is an essential assembly intermediate, from where, adding nucleotide (ADP is sufficient) could trigger their complete ring assembly. Still, there might be unknown assembly factors assisting the micro-complex formation. Furthermore, we propose a subunit-pair exchange/association mechanism for TRiC assembly, and our findings are beneficial for future TRiC-related therapeutic strategy development.

## Materials and Methods

### TRiC Subunit Expression

The yeast TRiC subunit genes were synthesized by PCR, and inserted into the pET28a vector with a N terminus 6xHis tag. The plasmids with recombinant single TRiC subunit were transformed into *E. coli* Rosetta (DE3). The cells were grown at 37°C to an OD_600_ of 0.6-0.8. The expression of the single TRiC subunits was induced by the addition of 0.5 mM isopropyl 1-thio-β-D-galactopyranoside, and the cells were further cultured at 18°C for 18 hrs and the cells were harvested by centrifugation.

### Purification of CCT homo-oligomer and yTRiC

The cells were lysed in the buffer with 20 mM HEPES • KOH pH 7.4, 300 mM NaCl, 10 mM MgCl_2_, 7 mM β-mercaptoethanol, 5% glycerol. One EDTA-free complete protease inhibitors EDTA free (Roche) was added in 100 mL lysis buffer. Specially, for CCT5, 1 mM ATP was added in the lysis buffer. The lysate was centrifuged at 20,000 g for 45 minutes, then the supernatant was passed over Ni-Beads (EMD Millipore Corporation). CCT single subunit was eluted off with 400 mM imidazole, and the fractions containing the CCT single subunit were combined and concentrated by Amicon Ultra 100 kDa MWCO centrifugal filter (Millipore). The concentrated CCT single subunit protein was passed over a Superose 6 10/300 GL size exclusion column (GE Healthcare). The individual subunit homo-oligomeric rings or micro-complexes exist around 11–14 ml fraction from the size exclusion column.

The yTRiC was purified following the published protocol (Zang et al., 2016). Briefly, the supernatant of yeast lysate was incubated with calmodulin resin (GE Healthcare) overnight at 4°C. Additionally, in the elution buffer, the original 10 mM EDTA was replaced by 2 mM EGTA.

### Recombination of dissociated CCT5 or yTRiC

Disassociated CCT5 purified without adding ATP throughout were incubated with 2 mM ATP or 2 mM ADP, respectively, for 1h min at room temperature (RT). We disassociated yTRiC complex by putting it in the relatively high-salt buffer without glycerol (300 mM NaCl, 10 mM MgCl_2_, 20 mM HEPES pH 7.4) at room temperature for 4h. The TRiC complexes were disassociated confirmed by negative-staining electron microscopy (NS-EM) (Fig. S8C). The disassociated TRiC were then incubated with 2 mM ATP or 2 mM ADP for 1h at RT, respectively, for it to reassemble.

### ATPase assay

The ATP-hydrolysis rates of CCT1-HR, CCT2-HR, CCT5-HSR-ATP, CCT6-HR, and WT TRiC were measured by performing a nicotineamide adenine dinucleotide (reduced) (NADH) coupled assay (Norby, 1988). All of the assays were conducted at room temperature in a buffer containing 20 mM Hepes/NaOH pH 7.4, 300 mM NaCl, and 10 mM MgCl_2_, and in the presence of 1 mM ATP. Experiments were performed in triplicate using 0.25 μM of the protein complex. Absorbance was measured in a 200 µl reaction volume using a 96-well plate reader tested by Synergy NEO (BioTeK). Data analysis was performed using GraphPad Prism 5.

### Luciferase refolding Assay

0.025 mg/ml of luciferase (Sigma) was unfolded in unfolding buffer (8 M guanidine hydrochloride, 20 mM HEPES • KOH pH 7.4, 2 mM DTT) at room temperature for 30 mins. The unfolded luciferase was diluted 100-fold into refolding buffer (20 mM Hepes pH 7.4, 50 mM NaCl, 2 mM DTT, 1 mM ATP, 10 mM MgCl_2_) with or without 0.2 mM purified HRs or yTRiC. At various time point, 5 μL refolding reaction was diluted 1:5 into luciferase assay system mix (Promega), and luminescence was measured on the GloMax-20/20 Luminometer (Promega). Data analysis was performed using GraphPad Prism 5.

### Thermal shift assay (TSA)

The thermal stability of the TRiC-like HRs and WT yTRiC was evaluated by thermal shift assay (TSA) (Liu et al., 2016). Briefly, all HRs and yTRiC were diluted into 0.5 mg/mL. Then, 5000× SYPRO Orange (Invitrogen, Carlsbad, CA, USA) was added to the reactions at a final concentration of 5×. The SYPRO orange binds the hydrophobic region and the hydrophobic region of the complexes can be exposed by thermal denaturation. All reactions were performed in triplicate in 384-well plates with a final volume of 10 μL. The thermal melting curve were monitored using a LightCycler 480 II Real-Time PCR System (Roche Diagnostics, Rotkreuz, Switzerland) with a ramp rate of 1 °C at the temperature range from 25 °C to 98 °C. The melting temperatures (Tm) were calculated by fitting the sigmoidal melting curve to the Boltzmann equation using GraphPad Prism 5.

### Cryo sample preparation and cryo-EM data acquisition

Purified TRiC-like HRs was diluted to 1.4 mg/ml, and 2.0 μl sample was applied to a glow-discharged holey carbon grid (R1.2/1.3, 200 mesh, Quantifoil). The grid was blotted with Vitrobot Mark IV (Thermo Fisher Scientific) and then plunged into liquid ethane cooled by liquid nitrogen. To handle the preferred orientation problem usually associated with group II chaperonins, the grid was pretreated with polylysine (Ding et al., 2017; Ding et al., 2019; Lander et al., 2012). Moreover, to prepare the HR-ATP-AlFx sample, the purified HR was incubated in the buffer (containing 1 mM ATP, 5 mM MgCl_2_, 5 mM Al (NO_3_)_3_, 30 mM NaF) for 1 h at 30 °C prior freezing.

The movies were collected on a Titan Krios electron microscope operated at an accelerating voltage of 300 kV, with a nominal magnification of 18,000x (yielding a pixel size of 1.318 Å after 2 times binning, Supplementary Table 1). The movie stacks were recorded on a Gatan K2 Summit direct electron detector operated in the Super resolution mode, with the defocus ranging from −0.8 to −2.5 μm. Each frame was exposed for 0.2 s, with an accumulation time of 7.6 s for each movie, leading to a total accumulated dose of 38 e^-^/Å^2^ on the specimen.

### Image processing and 3D reconstruction

We listed the image processing and reconstruction processes in Supplementary Table 1. Unless otherwise described, the data processing was performed in Relion 3.1 (Scheres, 2015). For all datasets, we performed the motion correction of each image stack using the embedded module of Motioncor2 (Zheng et al., 2017) in Relion 3.1, and determined CTF parameters by CTFFIND4 before further image processing (Mindell and Grigorieff, 2003). We picked ∼1000 particles to obtain several good class averages, which was used as template for further automatic particle picking; then excluded bad particles and ice contaminations by manual selection and 2D classification.

Take CCT2-HR-ATP-AlFx as an example (Fig. S4B), we performed multiple rounds of 3D classification to further clean up the particles, and obtained a dataset of 132,269 particles. We then carried out refinement without imposing symmetry in Relion 3.1 and obtained a map at 4.3-Å-resolution. Our rotational correlation analysis of the symmetry-free map using EMAN1.9 shows eight peaks with 45° interval, suggesting the existence of an 8-fold symmetry of the map. Thus, we imposed C8 symmetry in further refinement. After CTF refinement and Bayesian polishing, the CCT2-HR-ATP-AlFx map was refined to 3.2-Å-resolution (Fig. S4C-D). For CCT5^ΔNT^-HR-ATP, CCT6-HR and CCT6-HR-ATP-AlFx dataset, similar procedures were adopted as that used for the CCT2-HR-ATP-AlFx. Particularly, for tilted data of CCT6-HR and CCT6-HR-ATP-AlFx, we used goCTF software (Su, 2019) to determine the defocus for each of the particle, and we re-extracted these particles with corrected defocus and combined them with non-tilted data for further analysis. The final resolution for CCT5^ΔNT^-HR-ATP, CCT6-HR, and CCT6-HR-ATP-AlFx were estimated to be 3.7-, 5.9-, and 4.2-Å-resolution, respectively. For CCT1-HR and CCT1-HR-ATP-AlFx dataset, we performed multiple rounds of heterogeneous refinement in cryoSPARC (Punjani et al., 2017). Subsequently, after refinement in Relion 3.1 we carried out CTF refinement and Bayesion polishing, then the polished particles were non-uniform refined in cryoSPARC to 6.1- and 4.1-Å-resolutions for CCT1-HR and CCT1-HR-ATP-AlFx, respectively (Fig. S2, S3). For CCT2-HR and CCT5-HSR-ATP dataset, we performed multiple rounds of 3D classification; then for the best class, carried out CTF refinement and Bayesion polishing after refinement. Eventually, the polished particles were non-uniform refined in cryoSPARC to a resolution of 4.5 and 4.0 Å for CCT2-HR and CCT5-HSR-ATP, respectively. For all the reconstructions, the resolution estimations were accessed based on the gold-standard Fourier Shell Correlation (FSC) 0.143 criterion, and the local resolution was estimated using Resmap (Kucukelbir et al., 2014). UCSF Chimera and ChimeraX were used for map segmentation and figure generation (Goddard et al., 2018; Pettersen et al., 2004).

### Model building by flexible fitting

For CCT1-HR, CCT1-HR-ATP-AlFx, CCT2-HR, CCT6-HR, and CCT6-HR-ATP-AlFx structures, we used the model of the corresponding subunit from our previous yTRiC-NPP structure (PDB: 5GW4) as the initial model (Zang et al., 2016); for CCT2-HR-ATP-AlFx, we used the model of CCT2 from our previous TRiC-AMP-PNP structure (PDB: 5GW5) as the initial mode (Zang et al., 2016); for CCT5-HSR-ATP and CCT5^ΔNT^-HR-ATP, we used the model of CCT5 from our previous TRiC-ATP-AlFx (PDB: 6KS6) structure as the initial model (Jin et al., 2019). We fitted the model of the corresponding subunit multiple times into the TRiC-like HR map as a rigid body using UCSF Chimera, which were then combined as a complete model of the HR complex (Pettersen et al., 2004). Subsequently, we refined each of the models against the corresponding cryo-EM map using Rosetta (DiMaio et al., 2015) then Phenix (Adams et al., 2010). The final pseudo atomic models were validated using the *Phenix.molprobity* command in Phenix.

## Acknowledgments

We thank W. Yin (Shanghai Institute of Materia Medica, CAS) for the technique help in the TSA experiment. We are grateful to the staff of the NCPSS Electron Microscopy facility, Database and Computing facility, and Protein Expression and Purification facility for instrument support or technical assistance. This work was supported by grants from the CAS Pilot Strategic Science and Technology Projects B (XDB37040103), the National Basic Research Program of China (2017YFA0503503), the NSFC (31670754, 31861143028, and 31872714), the CAS Major Science and Technology Infrastructure Open Research Projects, the Program of Shanghai Academic Research Leader (20XD1404200), and the CAS-Shanghai Science Research Center (CAS-SSRC-YH-2015-01, and DSS-WXJZ-2018-0002).

## Author contributions

Y.C., C.L., and H.W. designed the experiments, C.L., H.W., W.H. S.W., and C.S. purified proteins and performed biochemical analysis, C.L. and H.W. collected cryo-EM data with the involvement of F.W., C.L., H.W., W.H., S.W., X.H, Q.Z. and C.S. performed particle picking, C.L., H.W. and M.J. performed 3D reconstruction and model building, C.L. and Y.C. analyzed the structures and wrote the manuscript.

## Data availability

All data presented in this study are available within the figures and in the Supplementary Information. Cryo-EM maps have been deposited in the EMDB with the accession numbers of EMD-***, and the associated models have been deposited in the Protein Data Bank with the accession numbers of ***.

The authors declare no conflict of interest.

## Supplementary Materials

**Fig. S1.**
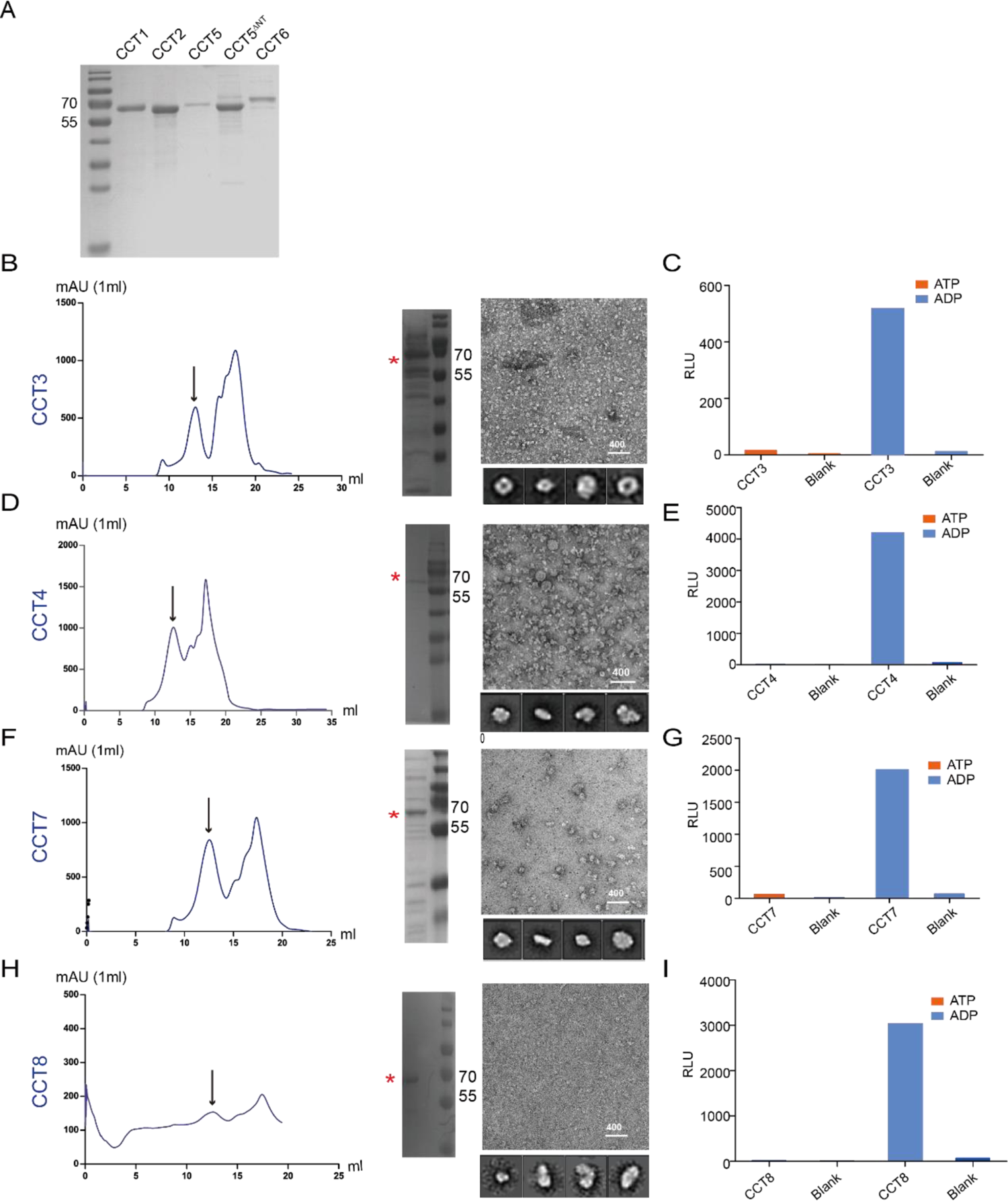
Purification and characterization of the yTRiC subunits expressed in *E. coli*. (A) SDS-PAGE of the subunits that could form ring-shaped structure (including CCT1, CCT2, CCT5-ATP, CCT5^ΔNT^-ATP, and CCT6), stained with Coomassie Brilliant Blue. (B, D, F, H) Gel-filtration graph, SDS-PAGE result, representative NS-EM micrograph, and reference-free 2D class averages for CCT3 (B), CCT4 (D), CCT7 (F), and CCT8 (H), respectively. The first peak in gel-filtration graph marked by black arrow corresponding to micro-complexes. In SDS-PAGE, CCT3/4/7/8 are indicated by red asterisk. (C, E, G, I) The ADP/ATP ratio assay result of CCT3 (C), CCT4 (E), CCT7 (G), and CCT8 (I) purified without added nucleotides, respectively, showing that CCT3/4/7/8 could load ADP from the environment of cells.

**Fig. S2.**
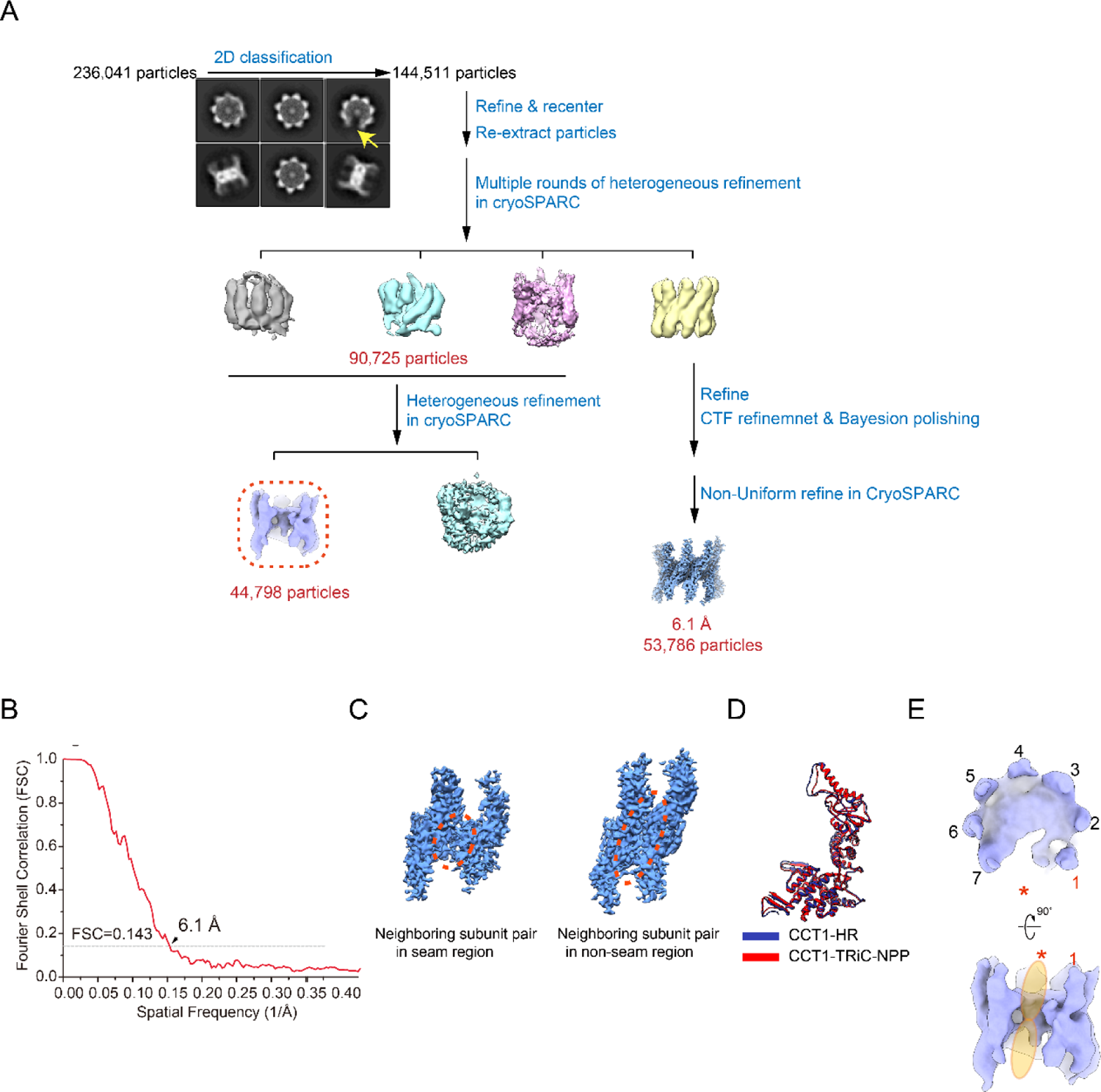
Cryo-EM data processing of CCT1-HR. (A) Cryo-EM data processing procedure of CCT1-HR dataset. Reference-free 2D class averages of CCT1-HR are also shown. The top view class average with a missing subunit-pair was indicated by yellow arrow. (B) Resolution assessment of CCT1-HR cryo-EM map by Fourier shell correlation (FSC) at 0.143 criterion. (C) The segmented map of the seam region (left) and the non-seam region (right) of CCT1-HR, illustrating the reduced interaction interface area between the two seam-subunit-pairs. (D) Model alignment of the CCT1 subunit from CCT1-HR (blue) and TRiC-NPP (red). (E) The cryo-EM map of the conformation with the seam-subunit-pair 8 (indicated by red asterisk) missing.

**Fig. S3.**
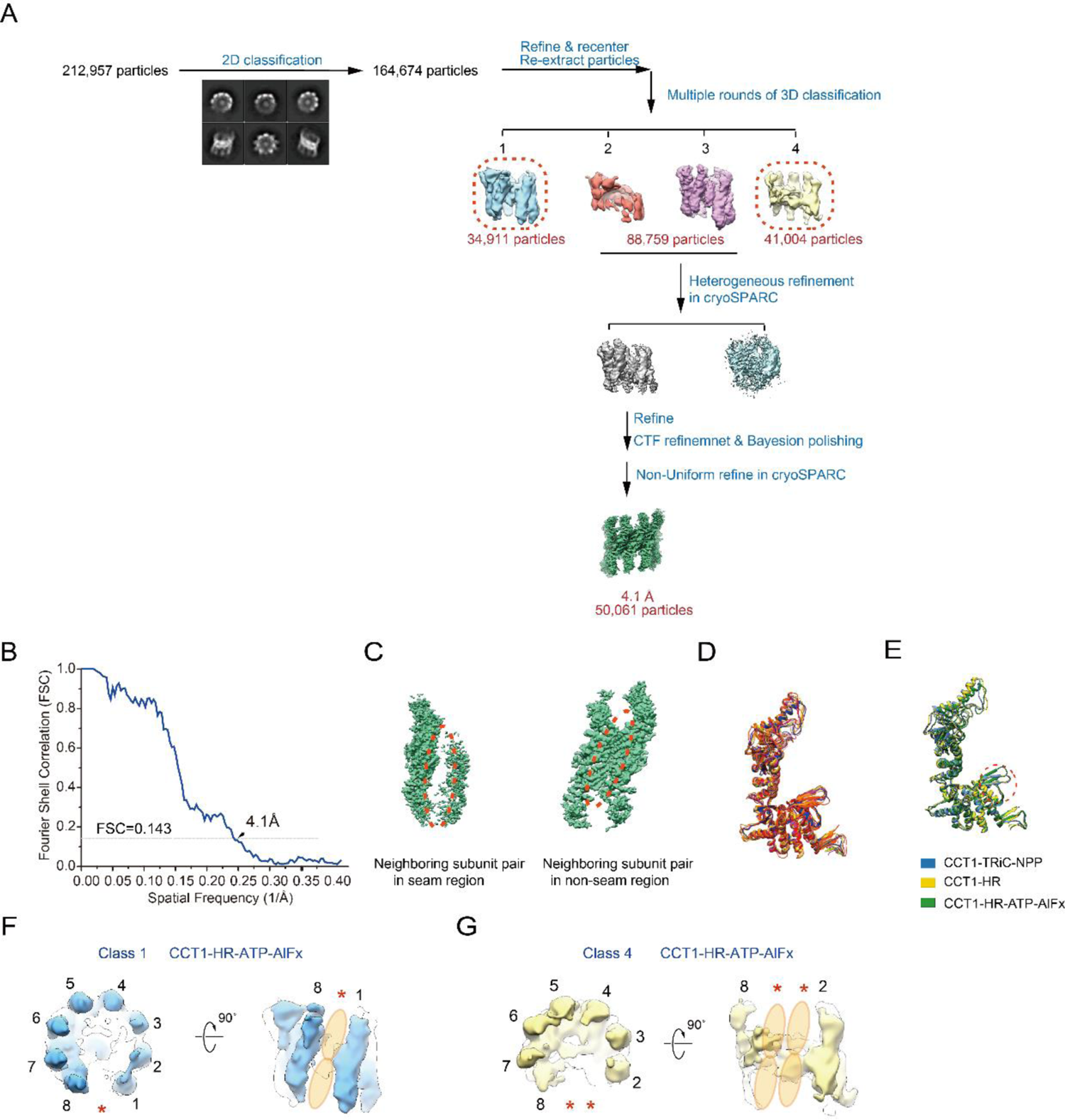
Cryo-EM data processing of CCT1-HR-ATP-AlFx. (A) Cryo-EM data processing procedure of CCT1-HR-ATP-AlFx dataset. Reference-free 2D class averages are also presented. The Class 1 & 4 with one or two missing subunit-pairs are indicated in the dotted red frame. (B) Resolution assessment of the CCT1-HR-ATP-AlFx map by the FSC at 0.143 criterion. (C) The map segmentation for the seam (left) and the non-seam (right) region of CCT1-HR-ATP-AlFx, illustrating the dramatically reduced interaction interface area between the two seam-subunit-pairs. (D) Alignment of the nine subunits of CCT1-HR-ATP-AlFx. (E) CCT1 subunit conformational comparison among TRiC-NPP, CCT1-HR, and CCT1-HR-ATP-AlFx. (F and G) The cryo-EM maps of Class 1 & 4, with one or two subunit-pairs missing in the seam region, respectively.

**Fig. S4.**
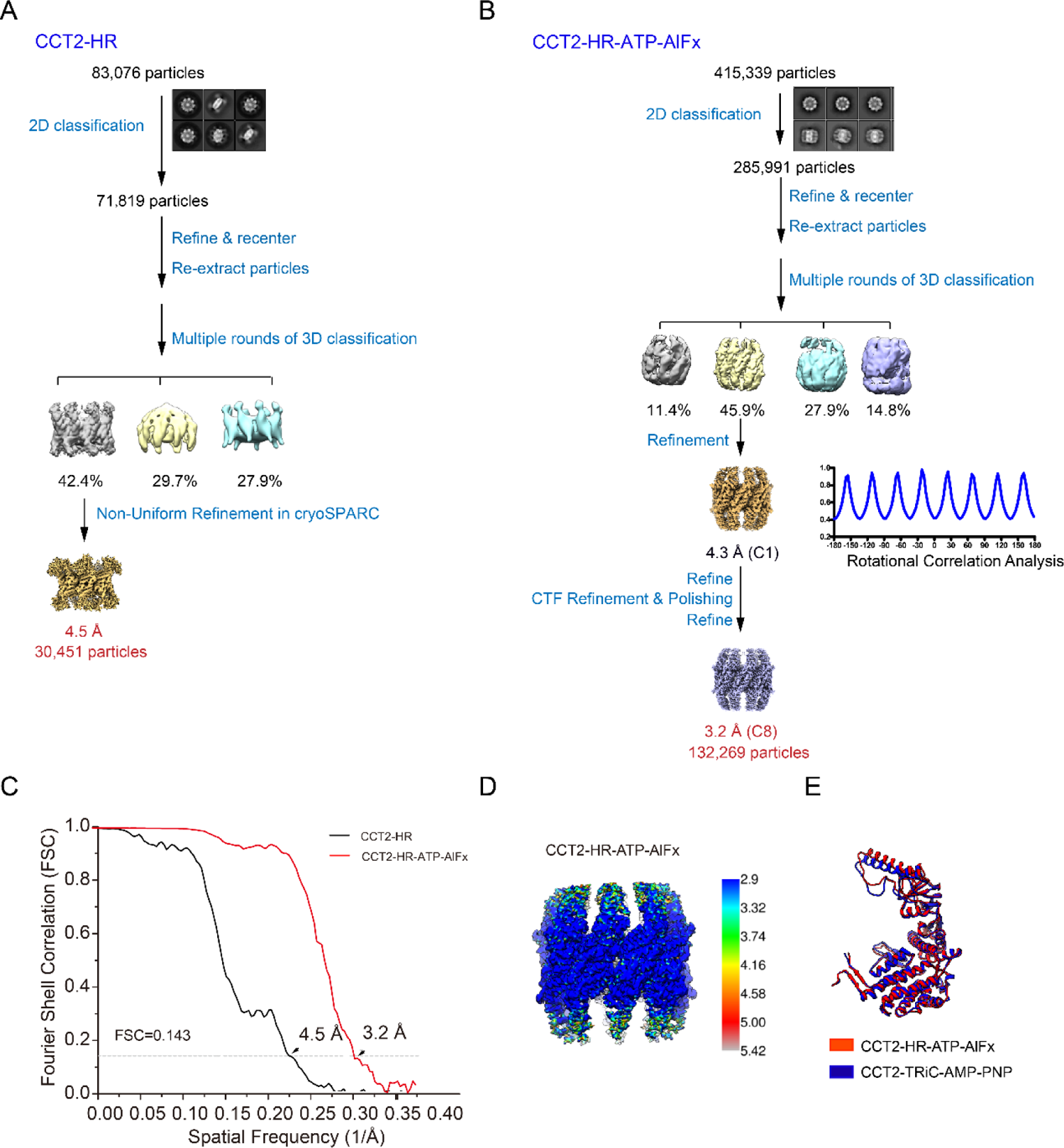
Cryo-EM data processing of CCT2-HR and CCT2-HR-ATP-AlFx. (A-B) Data processing procedure of CCT2-HR (A) and CCT2-HR-ATP-AlFx (B). For CCT2-HR-ATP-AlFx, the rotational correlation analysis of the map without imposing any symmetry (C1 map) shows eight peaks with 45° interval, suggesting the existence of an 8-fold symmetry in the map. (C) Resolution assessment of the CCT2-HR and CCT2-HR-ATP-AlFx maps by the FSC curve at 0.143 criterion. (D) Local resolution estimation of the CCT2-HR-ATP-AlFx map by Resmap. (E) Model alignment of the CCT2 subunits from the CCT2-HR-ATP-AlFx (red) and TRiC-AMP-PNP (PDB: 5GW5, blue) structures, indicating no obvious conformational variations between them.

**Fig. S5.**
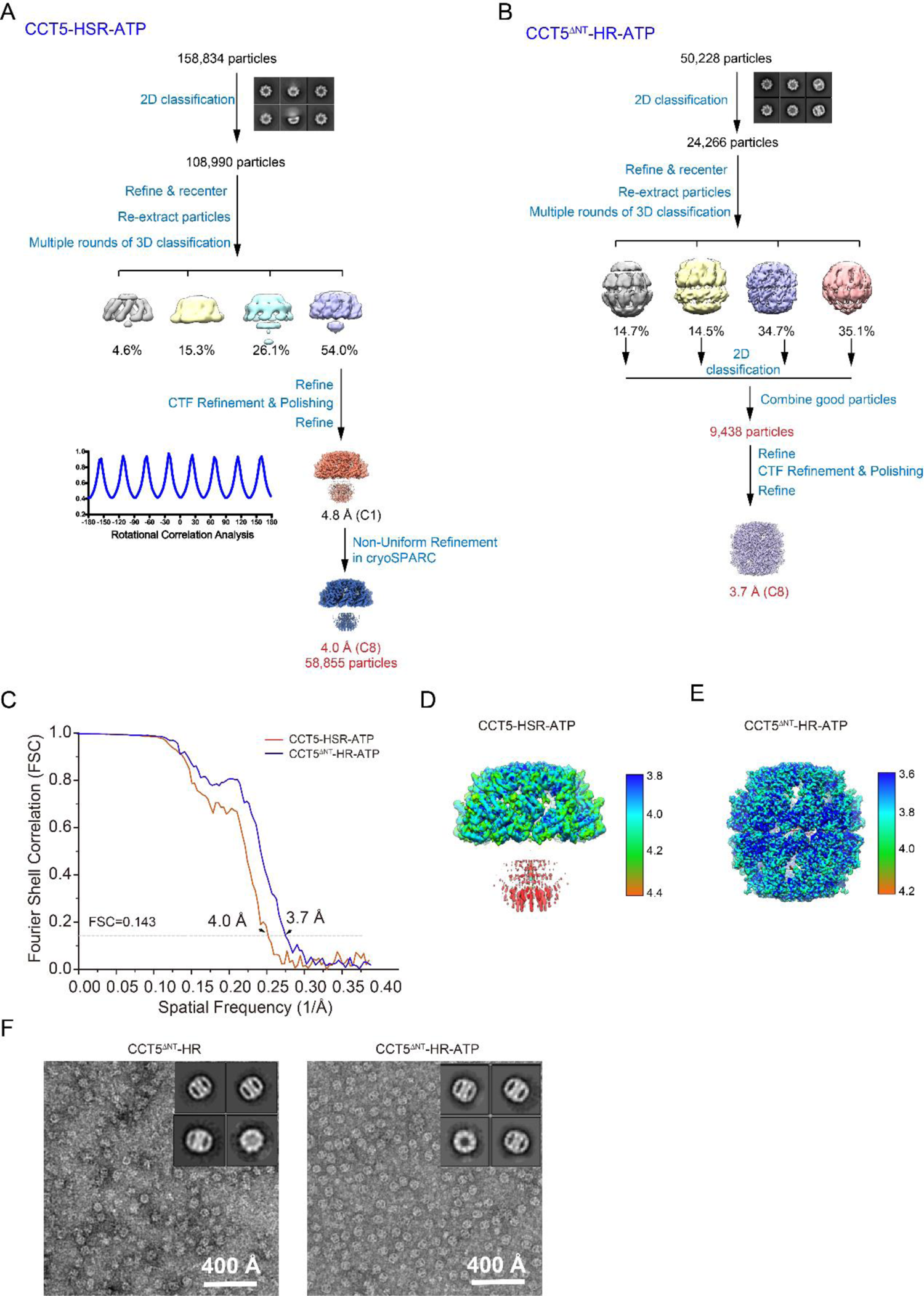
Cryo-EM data processing of CCT5-HSR-ATP and CCT5^ΔNT^-HR-ATP. **(A-**B) Cryo-EM data processing procedure for CCT5-HSR-ATP and CCT5^ΔNT^-HR-ATP. The rotational correlation analysis for CCT5-HSR-ATP indicating the existence of an 8-fold symmetry in the structure. (C) Resolution assessment of the CCT5-HSR-ATP and CCT5^ΔNT^-HR-ATP maps by the FSC curve at 0.143 criterion. (D-E) Local resolution estimations of the CCT5-HSR-ATP (D) and CCT5^ΔNT^-HR-ATP maps (E) by Resmap. (F) The representative negative staining micrographs and reference-free 2D averages of CCT5^ΔNT^-HR (left) and CCT5^ΔNT^-HR-ATP (right), respectively.

**Fig. S6.**
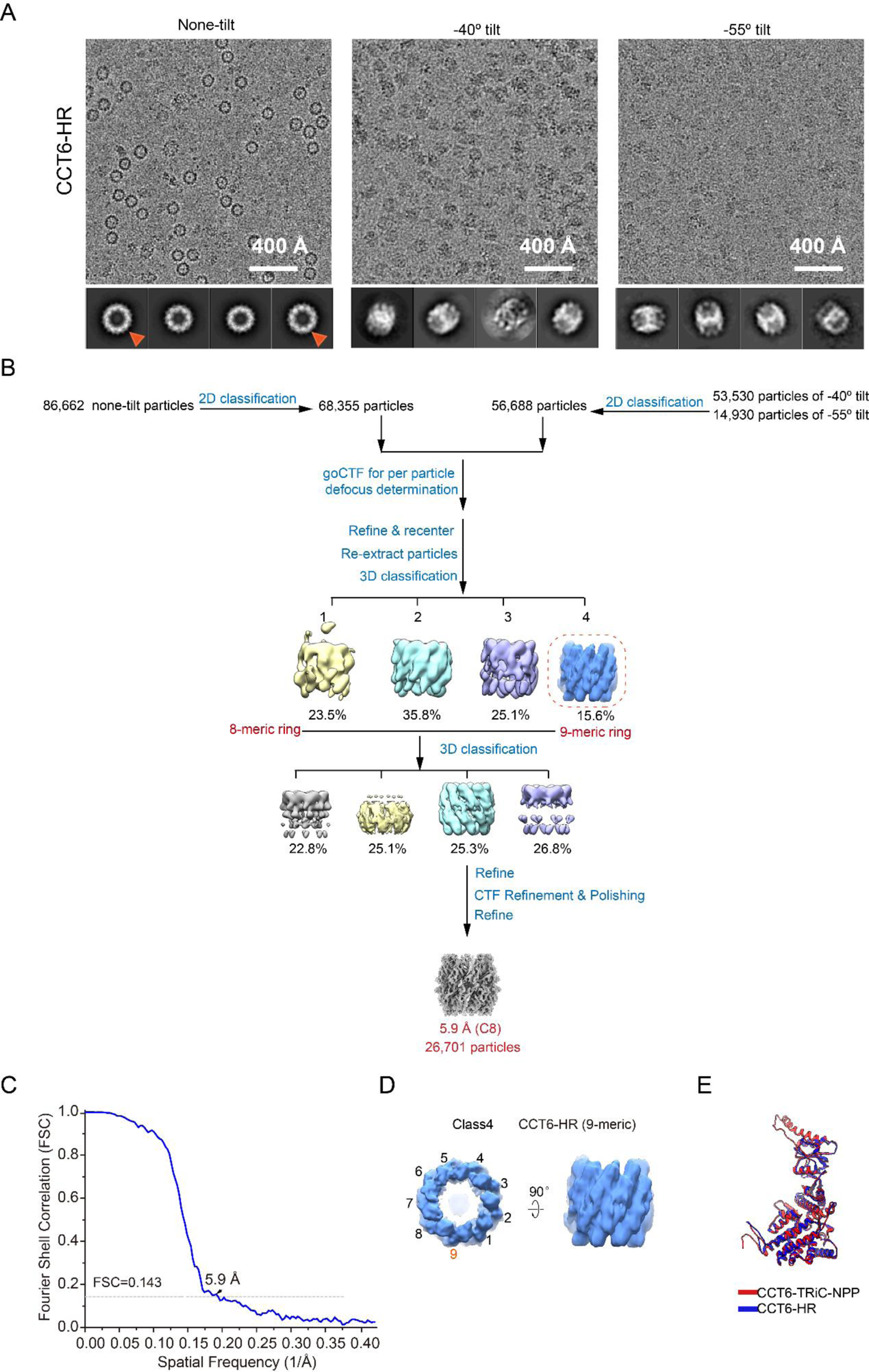
Cryo-EM data processing of CCT6-HR. (A) Representative cryo-EM micrographs of CCT6-HR collected at 0°, −40° and −55° tilt, respectively, and the corresponding reference-free 2D class averages. The top view 2D class averages at larger diameter are indicated by orange arrow head. (B) Cryo-EM data processing procedure of CCT6-HR. The 9-meric CCT6-HR is indicated by a dotted red frame. (C) Resolution assessment of the 8-meric CCT6-HR cryo-EM map by the FSC curve at 0.143 criterion. (D) The Class 4 reconstruction represents the 9-meric CCT6-HR. (E) Model alignment of the CCT6 subunit from CCT6-HR (blue) and TRiC-NPP (red), indicating no conformational variations between them.

**Fig. S7.**
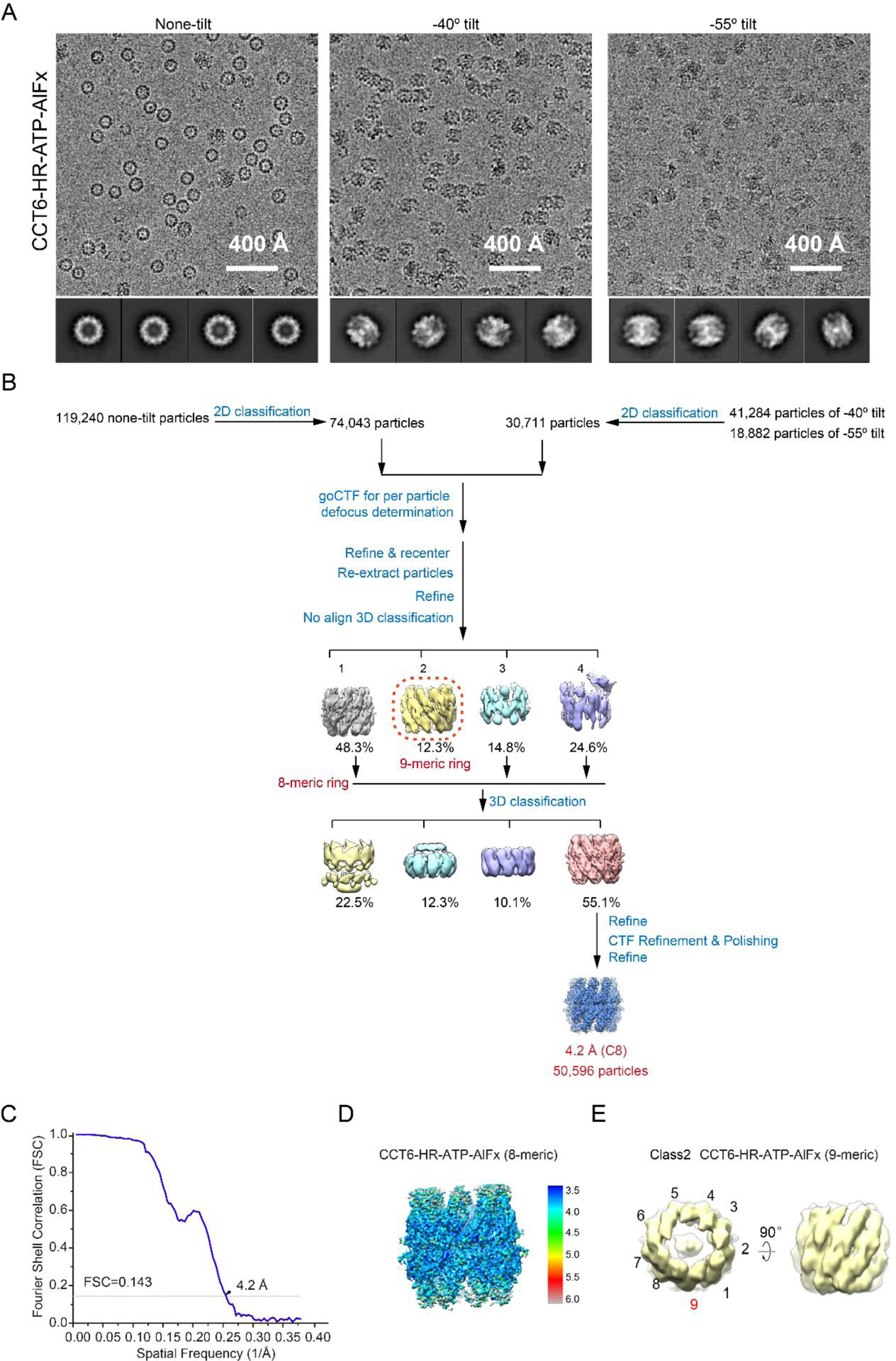
Cryo-EM data processing of CCT6-HR-ATP-AlFx. (A) Representative cryo-EM micrographs of CCT6-HR-ATP-AlFx collected at 0°, −40° and −55° tilt, respectively, and the corresponding reference-free 2D class averages. (B) Cryo-EM data processing procedure of CCT6-HR-ATP-AlFx. The 9-meric CCT6-HR-ATP-AlFx is indicated by dotted red frame. (C) Resolution assessment of the 8-meric CCT6-HR-ATP-AlFx cryo-EM map by the FSC curve at 0.143 criterion. (D) Local resolution estimation of the CCT6-HR-ATP-AlFx map by Resmap. (E) The Class 2 reconstruction represents the 9-meric CCT6-HR-ATP-AlFx.

**Fig. S8.**
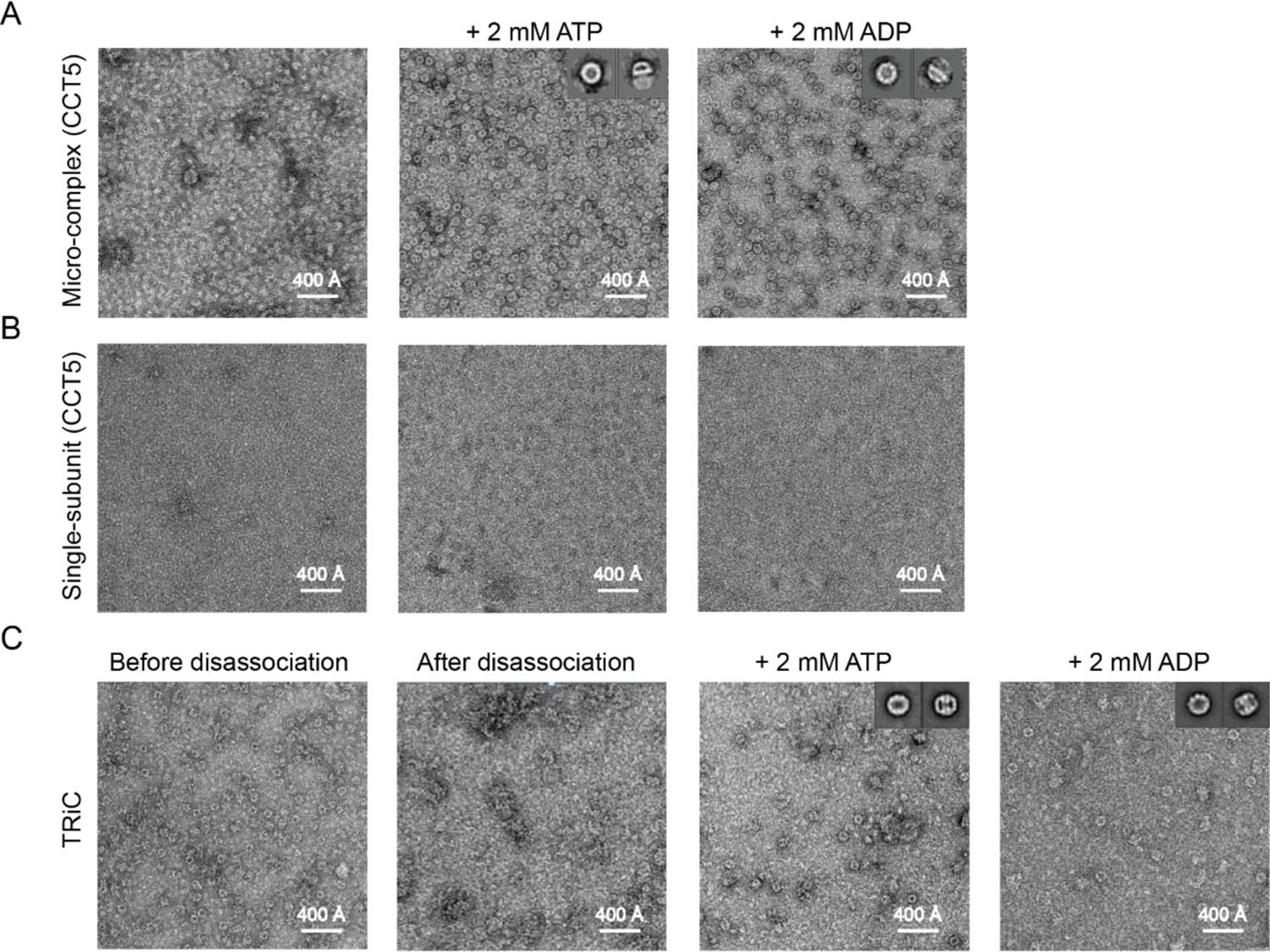
Nucleotide plays an important role in TRiC ring assembly. (A and B) CCT5 micro-complex can re-assemble into TRiC-like ring structure in the presence of either ATP or ADP (A), while CCT5 single subunit cannot (B). (C) Representative negative staining micrograph of TRiC before dissociation (left) or being dissociated (right). (D) Dissociated TRiC can re-assemble into rings in the presence of 2 mM ATP (close state) or ADP (open state), indicating the ADP is sufficient to trigger the intermediate state micro-complex to re-assemble into intact TRiC complex, while ATP could trigger the re-assembled TRiC to close the ring.

**Table S1.**
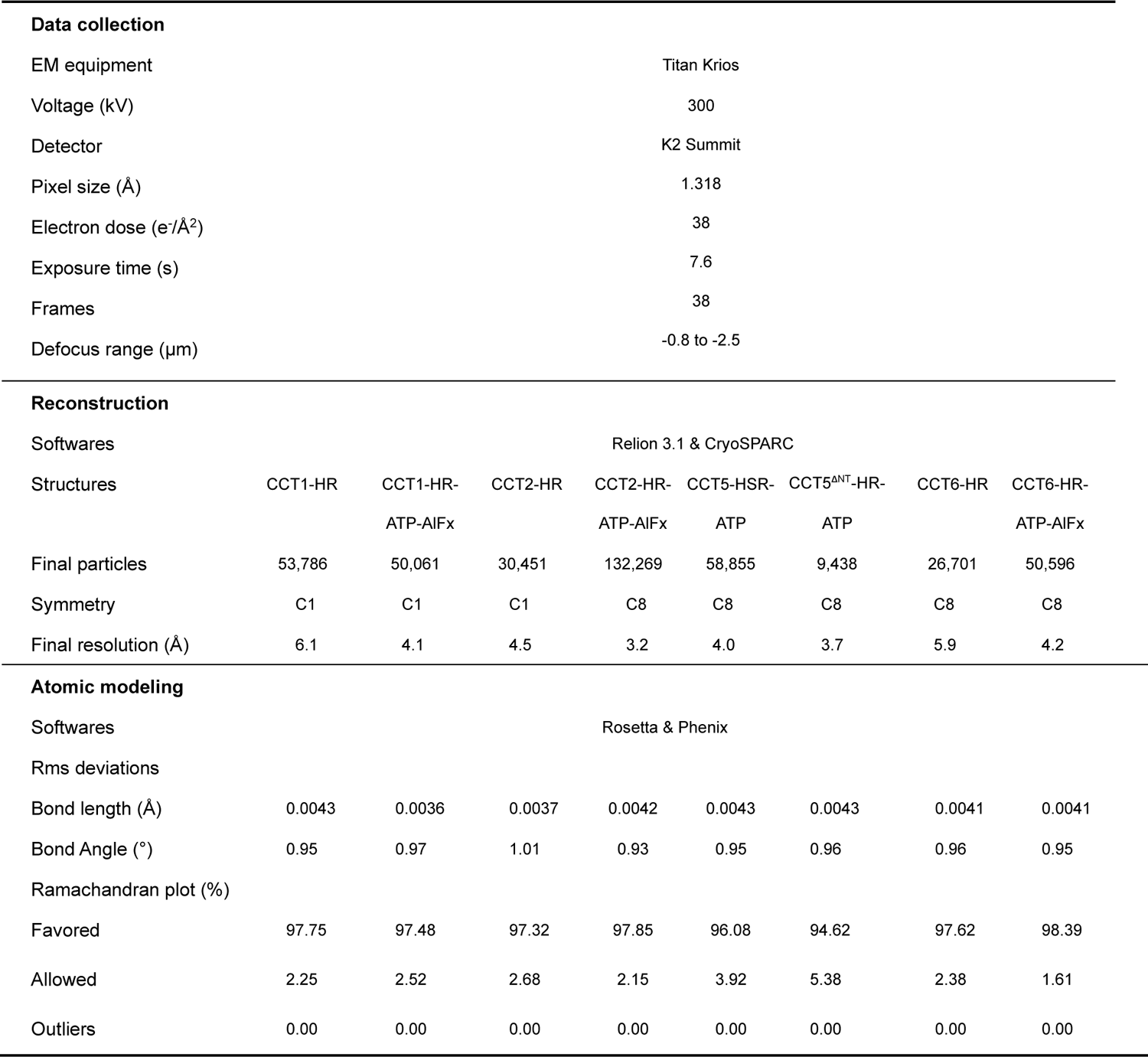
Statistics of cryo-EM data collection and processing

